# Muscle Cathepsin B treatment improves behavioral and neurogenic deficits in a mouse model of Alzheimer’s Disease

**DOI:** 10.1101/2025.01.20.633414

**Authors:** Alejandro Pinto, Hazal Haytural, Cássio Morais Loss, Claudia Alvarez, Asude Ertas, Olivia Curtis, Alyssa R. Williams, Grayson Murphy, Ken Salleng, Sylvia Gografe, Ali Altıntaş, Tal Kafri, Romain Barres, Atul S. Deshmukh, Henriette van Praag

## Abstract

Muscle secretes factors during exercise that enhance cognition. Myokine Cathepsin B (Ctsb) is linked to memory function, but its role in neurodegenerative disease is unclear. Here we show that AAV-vector-mediated Ctsb overexpression in skeletal muscle in an Alzheimer’s Disease (AD) mouse model (APP/PS1), improves motor coordination, memory function and adult hippocampal neurogenesis, while plaque pathology and neuroinflammation remain unchanged. Additionally, in AD mice, Ctsb treatment modifies hippocampal, muscle and plasma proteomic profiles to resemble that of wildtype controls. Conversely, in wildtype mice, Ctsb expression causes memory deficits and results in protein profiles across tissues that are comparable to AD control mice. In AD mice, Ctsb treatment increases the abundance of hippocampal proteins involved in mRNA metabolism and protein synthesis, including those relevant to adult hippocampal neurogenesis and memory function. Furthermore, Ctsb treatment enhances plasma metabolic and mitochondrial processes, and reduces inflammatory responses. In muscle, Ctsb expression elevates protein translation in AD mice, whereas in wildtype mice mitochondrial proteins decrease. Overall, the biological changes in the treatment groups are consistent with effects on memory function. Thus, skeletal muscle Ctsb application has potential as an AD therapeutic intervention.

## INTRODUCTION

Alzheimer’s Disease (AD) is a prevalent neurodegenerative disease[1], with minimal and largely ineffective pharmacological treatment options[2]. Brain pathology includes amyloid plaques, tau protein aggregation, neuroinflammation, synaptic dysfunction and neural tissue loss[3]. Lifestyle plays an important role in dementia incidence. Sedentary behavior and loss of muscle mass and strength (sarcopenia) is linked to hippocampal dysfunction, AD risk and progression[4-7]. Conversely, physical activity enhances brain volume, connectivity, blood flow, cognition and mood, and may delay or prevent onset of the disease[8-10]. In rodents, running increases adult hippocampal neurogenesis, synaptic plasticity, neurotrophins and memory function, and reduces neuroinflammation and anxiety. Research into the underlying mechanisms indicates that peripheral factors produced during exercise can convey neurogenic and neuroprotective effects[11, 12]. Infusions of blood derived from young mice or exercising donors decreases neuroinflammation, and enhances structural and functional neuroplasticity in aged animals[13, 14]. Factors secreted from muscle, liver, adipose tissue and platelets during exercise into circulation may mediate such effects[15-18] and hold promise as novel treatments for cognitive decline.

We identified Cathepsin B (Ctsb), a lysosomal cysteine protease[19], that has been primarily studied in pathological contexts - where it is linked to cancer progression, exacerbation of brain injury and equivocally in AD[20-23] -, as a novel myokine[24]. In humans Ctsb is upregulated in circulation by exercise [24-30] and electrical muscle stimulation[31], and is associated with cognition[24, 25, 32]. *Ctsb* ablation in mice precludes running-induced pro-neurogenic and mnemonic effects[24]. Whether this is mediated by muscle *Ctsb* loss is unclear. Running[24] and muscle-specific PGC-1α overexpression increase muscle *Ctsb* levels[33, 34]. In addition, constitutive overexpression of transcription factor E-B (TFEB), a master regulator of lysosomal function and cellular metabolism[35], enhances skeletal muscle *Ctsb* expression, reduces neuroinflammation and improves cognition in aged mice[36, 37]. Furthermore, in *Drosophila,* skeletal muscle transcriptional upregulation of proteases is neuroprotective in the brain and retina[17, 38]. However, it remains to be determined whether muscle Ctsb treatment can prevent neurogenic and cognitive decline in neurodegenerative disease.

Here we explored effects of skeletal muscle Ctsb expression in an AD mouse model, expressing chimeric mouse/human Swedish amyloid precursor protein (APP) and presenilin delta 9 mutations (APP/PS1), that cause neuropathological changes such as amyloid plaques, gliosis and cognitive deficits[39-41]. In AD mice, skeletal muscle Ctsb expression prevented the appearance of motor, cognitive and neurogenic deficits, and modified tissue and plasma proteomic profiles to levels of control WT mice. Our findings show that muscle Ctsb treatment protects against memory decline in AD mice.

## RESULTS

Four-month-old male mice (AD and WT mice) received AAV-vector-mediated Ctsb or Control (Con) treatment: WT-Con, WT-Ctsb, AD-Con, and AD-Ctsb. Six months later, mice underwent a battery of behavioral tests, consisting of activity box (open field), rotarod, Morris water maze, and fear conditioning (Figure 1A). Immunohistochemical analysis of adult-born neurons, markers of neuroinflammation (microglia and astrocytes), and AD-related pathology was assessed in the hippocampus and cortex. Furthermore, hippocampus, gastrocnemius muscle and plasma proteome were analyzed by liquid chromatography-mass spectrometry (LC-MS/MS) (Figure 1B).

**Fig 1.**
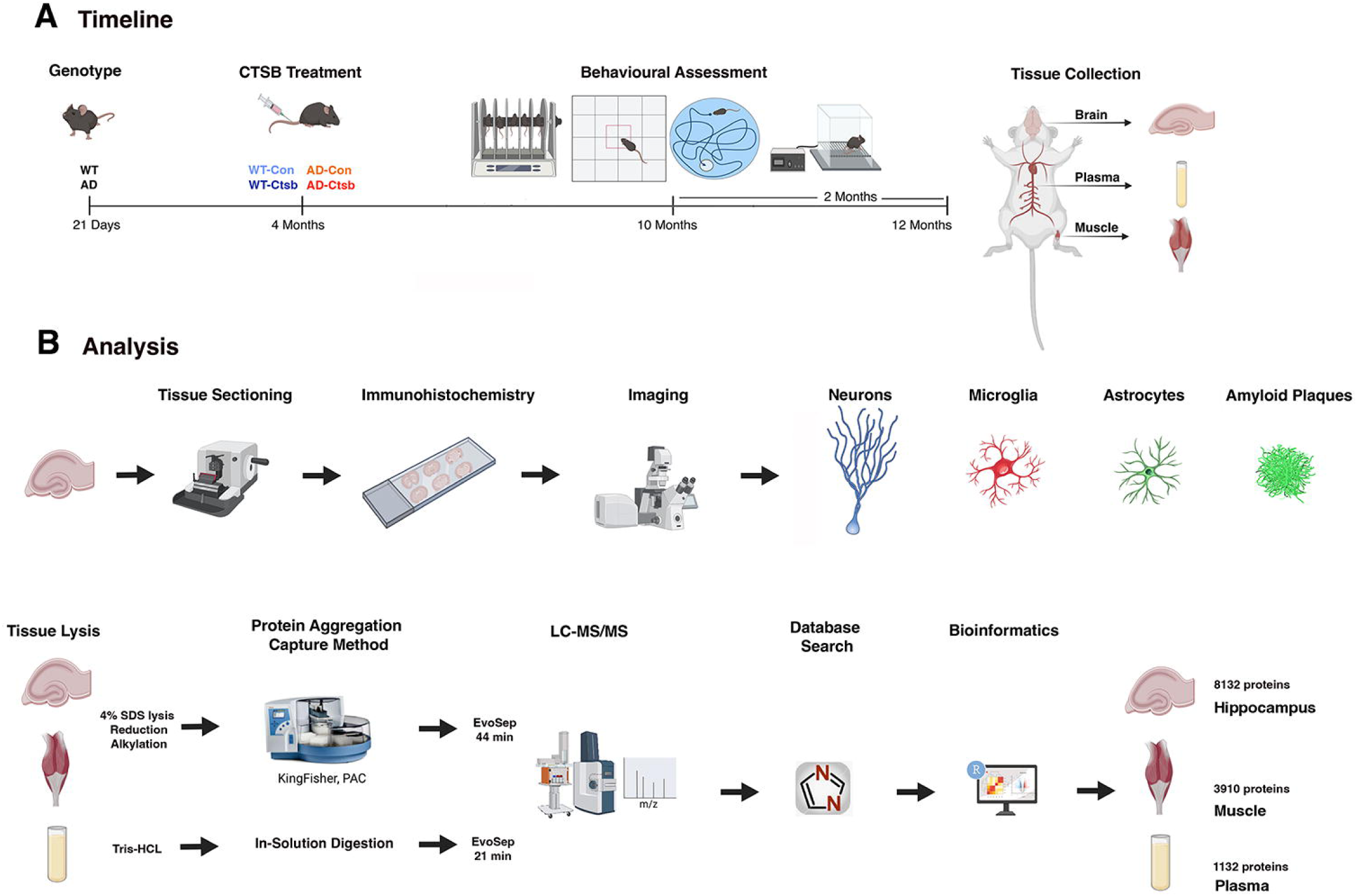
Experimental design and timeline. **(A)** Four-month-old male mice (either AD or WT mice) were injected into the tail vein with Ctsb (pAAV9-tMCK-mCTSB-IRES-eGFP) or Control (Con) vector (pAAV9-tMCK-IRES-eGFP): WT-Con (N=10), WT-Ctsb (N=9), AD-Con (N=8), and AD-Ctsb (N=14). Mice were subjected to a battery of behavioral tests from 10 months of age. The behavioral assessment consisted of activity box (open field), rotarod, Morris water maze, and fear conditioning paradigm. At twelve months of age mice were deeply anesthetized and euthanized for tissue and plasma collection. **(B)** One hemisphere of the brain tissue was sequentially sectioned and used for histological evaluation of adult neurogenesis, amyloid beta plaque deposition, and neuroinflammation. Hippocampus from the other brain hemisphere, gastrocnemius muscle and plasma were used for proteomic assays. Briefly, hippocampal and muscle tissues were lysed in 4% SDS buffer, following reduction and alkylation steps, protein aggregation capture method. Cleaned peptides were injected into LC-MS/MS (Evosep HPLC 44 minutes coupled to timsTOF Pro 2 mass spectrometer). Meanwhile plasma proteins were solubilized directly in Tris buffer. Cleaned peptides were injected into LC-MS/MS (Evosep HPLC 21 minutes coupled to timsTOF Pro 2 mass spectrometer).

### Locomotor behavior is modified by Ctsb treatment

In order to assess the impact of AAV-vector-mediated skeletal muscle Ctsb expression on locomotor activity and motor coordination, mice were tested within the activity box (open field) and rotarod apparatus.

#### Activity Box and Rotarod

Mice were placed in the activity box (open field) for 60 minutes. The total ambulatory distance in WT-Ctsb but not AD mice was increased (Figure 2A). Analysis of distance traveled over time showed similar habituation to the arena between groups (Figure S1; Tables S1 and S2). In the rotarod the AD-Con group had a shorter latency to fall as compared to WT-Con and AD-Ctsb groups. Thus, Ctsb expression reversed the motor coordination deficit in AD animals (Figure 2B; Table S3).

**Fig 2.**
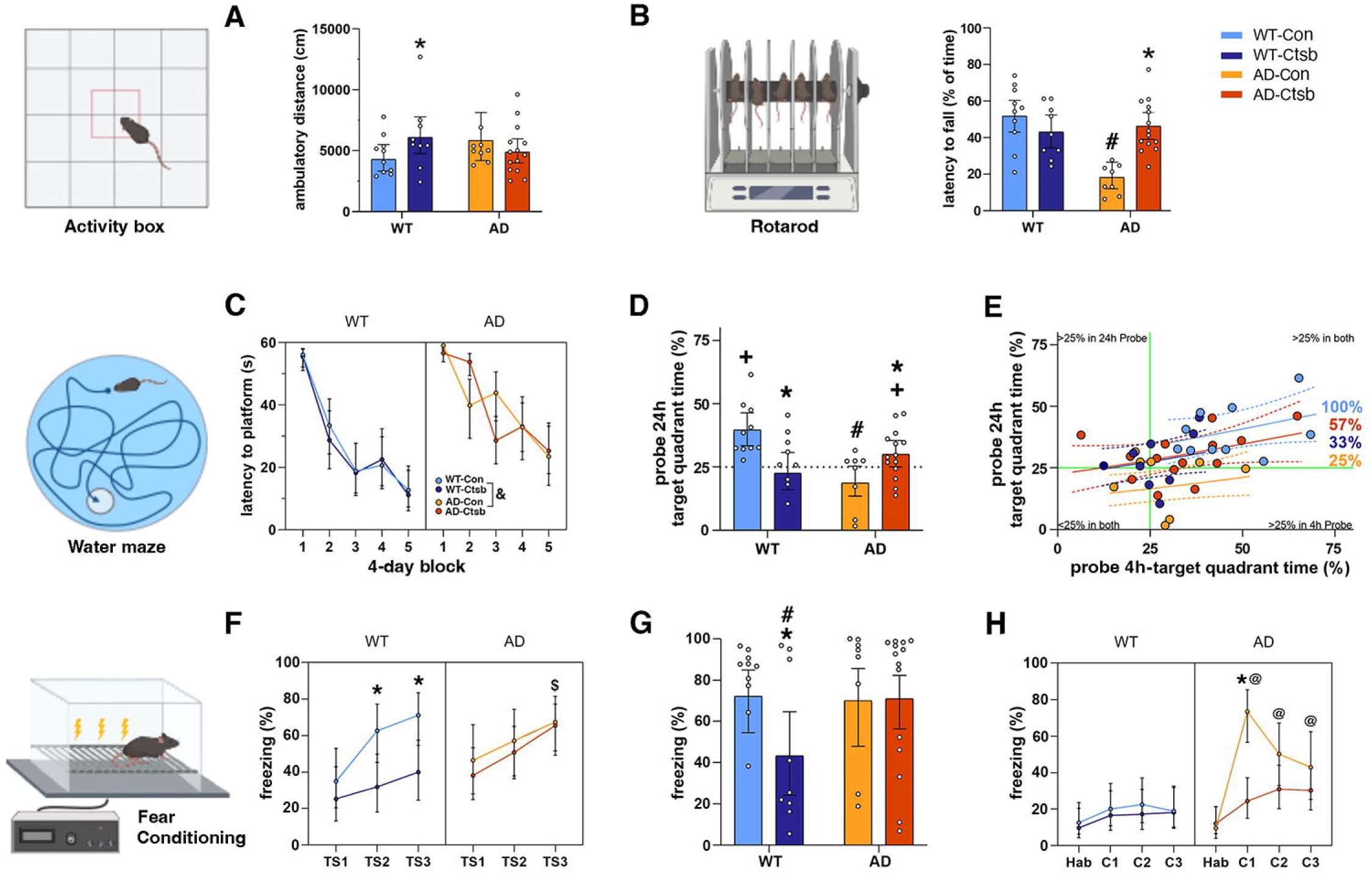
Ctsb treatment differentially affects locomotion, motor coordination and memory function in WT and AD mice. **(A)** Ctsb treatment increased distance traveled in WT mice during a 60 min activity box session. **(B)** AD-Con mice had a shorter latency to fall than all the other groups on an accelerating rotarod (4-40 rpm). **(C)** Acquisition curves in the Morris water maze, analyzed over 4-day bins (during a 20-day training protocol). WT mice perform better than AD mice throughout the acquisition phase. **(D, E)** Morris water maze probe trials to test retention of spatial memory. (**D**) AD-Con mice and WT-Ctsb mice had impaired retention of spatial memory performance during the 24h probe trial. Only WT-Con and AD-Ctsb mice preferred the platform quadrant compared to 25% chance. (**E**) Considering the performance in the 4h and 24h probe trials as a whole, WT-Con (100%), AD-Ctsb (57%), AD-Con (25%) and WT-Ctsb (33%) mice spent more than 25% of time in the target quadrant in both trials. Solid green lines represent 25% chance. **(F)** In the fear conditioning paradigm WT-Ctsb mice froze less during the conditioning session and **(G)** the tone-cued phase, as compared to the other groups. (**H**) AD-Con mice displayed increased CFR to context as compared to the other groups. (N; WT-Con = 10, WT-Ctsb = 9, AD-Con = 8, AD-Ctsb = 14). ^#^P<0.05 AD compared to WT in the same Treatment. *P<0.05 Ctsb compared to Con in the same Genotype. ^+^P<0.05 of being equal or lesser than 25% chance (one-tailed Wilcoxon signed rank exact test). ^&^P<0.05 AD compared to WT (Genotype main effect). ^$^P<0.05 AD-Ctsb compared to WT-Ctsb. ^@^P<0.05 AD-Con compared to WT-Con. Data were analyzed by GLM **(A, B, D, G)** or GLMM **(C, F, H)** and are presented as Estimated Marginal Means and their respective 95% CI. In **E**, solid lines represent the Estimated Marginal Means while dashed lines represent 95% CI.

### Learning and Memory is differentially affected by Ctsb in AD versus WT mice

#### Morris water maze

To assess spatial learning, mice were trained in the Morris water maze[42] with 4 trials per day for twenty days. In the acquisition phase, the WT groups displayed a shorter latency to the platform than the AD groups (4-day bins over time, Figure 2C, and Table S4). For all groups, mean latency to the platform was <20 sec in the last training session. Swim speed did not differ between the groups on day 1 and did not change over time (Figure S2, Tables S5 and S6).

To test memory retention, the platform was removed and probe trials were conducted 4h (Figure S2) and 24h after the last training session (Tables S7, S8). In the 24h probe trial, the WT-Con and AD-Ctsb groups spent more time in the target quadrant than the AD-Con group. In addition, WT-Ctsb mice were in the target quadrant for a shorter time than WT-Con mice. Both the WT-Con and AD-Ctsb groups preferred the target quadrant above 25% chance (Figure 2D, Tables S9, S10), indicating that these groups remember where the platform was located during the acquisition phase. Examining the performance in the 4h and 24h probe trials as a whole, we found that 100% WT-Con, 57% AD-Ctsb, 33% WT-Ctsb and 25% AD-Con mice, spent more than 25% of time in the target quadrant in both trials (Figure 2E). Altogether, these results indicate that the intervention benefits spatial memory retention in AD mice, whereas it has a detrimental outcome in WT mice.

#### Fear Conditioning

Mice were subjected to a Pavlovian conditioning-based approach to assess fear learning and memory, by measuring the degree of freezing, a Conditioned Fear Reaction (CFR), elicited by pairing a tone with a shock (Figure 2F, 2G and 2H). In the habituation (Hab) session on Day 1, there were no differences in freezing between the groups (WT-Con = 12.66; 6.35 – 23.65; WT-Ctsb = 9.80; 4.37 – 20.50; AD-Con = 9.61; 4.02 – 21.29; AD-Ctsb = 12.12; 6.52 – 21.42, % freezing). On Day 2, the Conditioning Phase consisted of three tone-shock (TS) pairings. The WT-Con, AD-Con and AD-Ctsb mice presented higher CFR than WT-Ctsb mice. Indeed, WT-Ctsb mice freeze less (<40%) than all the other groups (> 65%) in Tone-Shock 3 (TS3), the last conditioning stimulus (CS) – unconditioned stimulus (US) presentation (Figure 2F; Table S11). On Day 3, mice were first tested for tone-cued memory in a new contextual environment (T1-T3). The WT-Ctsb group exhibited a reduced average tone-elicited CFR (<44% freezing) compared to the other groups (>70% freezing) (Figure 2G; Table S12). One hour after the tone-cued session on day 3 (C1), as well as on days 5 and 7 (C2 and C3, respectively), mice were tested in the original environment used in the habituation (Hab) and conditioning sessions. Elevated context-elicited CFR was observed in AD-Con only (Figure 2H; Table S13). Overall, fear conditioning behavior in AD-Ctsb mice was similar to that of the WT-Con group (Figure 2F, 2G and 2H).

### Adult Hippocampal Neurogenesis is enhanced by Ctsb in AD mice

To assess adult hippocampal neurogenesis at 12-months of age, we quantified doublecortin (DCX)^+^ cells (Figure 3A-D). Analysis showed an interaction between genotype and treatment for DCX, an endogenous immature neuron marker[43]. Specifically, DCX^+^ cell number in AD-Con was reduced by 50% as compared to the WT-Con group. Treatment with Ctsb restored DCX^+^ cell number to WT-Con levels in AD mice (Figure 3E, Table S14). Cell genesis at 5-months of age did not differ between groups (Figure S3, Table S15). Our results indicate that AAV-mediated Ctsb expression in skeletal muscle prevented the adult hippocampal neurogenesis deficit caused by AD.

**Fig 3.**
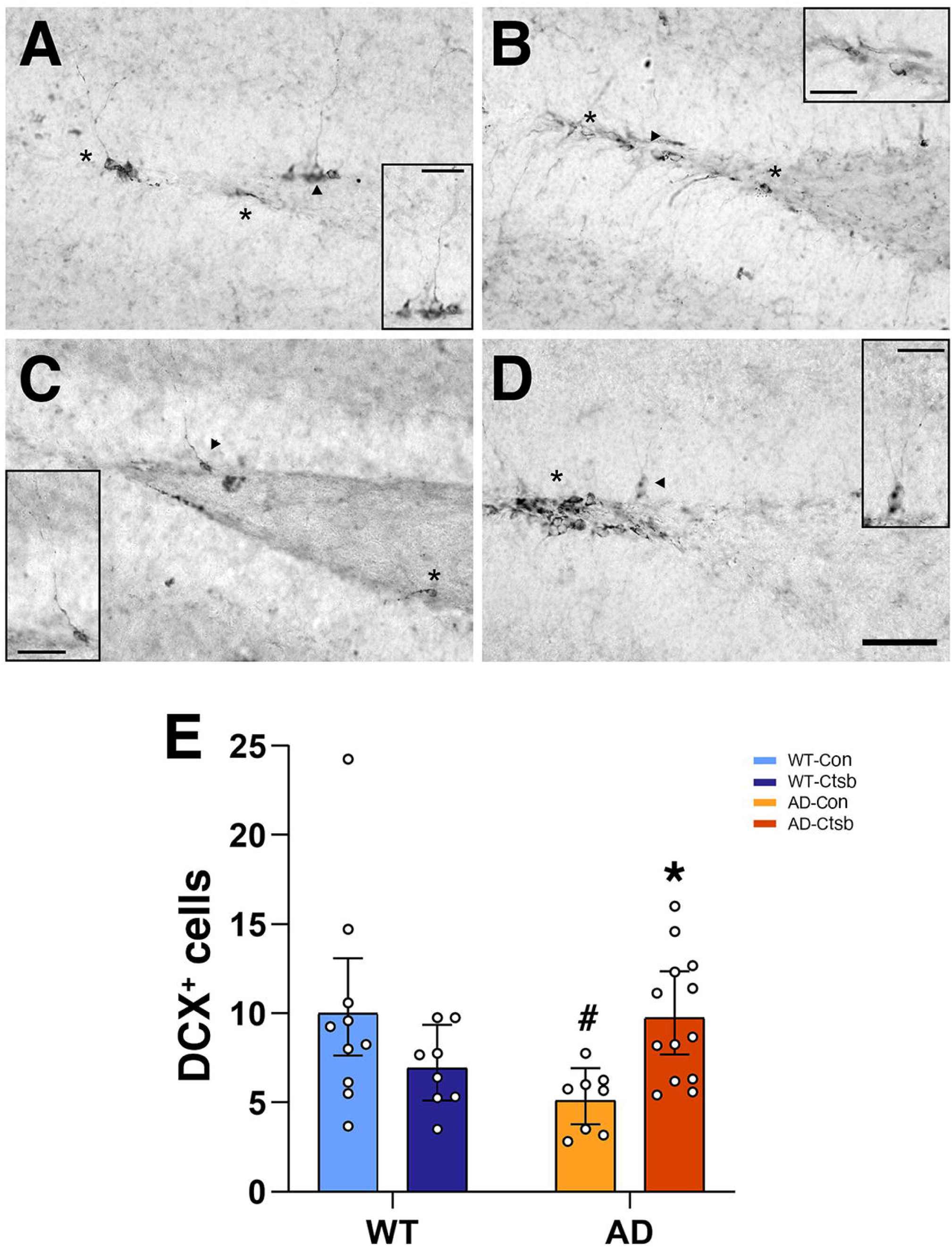
Ctsb treatment improves adult neurogenesis in AD mice. **(A-D)** Representative photomicrographs of the dentate gyrus derived from **(A)** WT-Con; **(B)** WT-Ctsb; **(C)** AD-Con; **(D)** AD-Ctsb brain tissue sections subjected to DCX staining. **(E)** Treatment with Ctsb in AD mice restored DCX cell number to WT-Con levels. (N; WT-Con = 10, WT-Ctsb = 8, AD-Con = 8, AD-Ctsb = 13). Scale bars: 50 μm overview; 25 μm inset. ^#^P<0.05 AD compared to WT in the same Treatment. *P<0.05 Ctsb compared to Con in the same Genotype. Data in **E** was analyzed by GLM and are presented as Estimated Marginal Means and their respective 95% CI.

### Plaque deposition and microglial activation in AD mice

To evaluate AD pathology, sections derived from AD-Con and AD-Ctsb mice were stained with Thio-S to label amyloid deposits (Figure 4A,B). Plaque density (Figure 4C,E) and counts (Figure 4D,F) in AD cortex and hippocampus did not differ between groups (Tables S17-S20).

**Fig 4.**
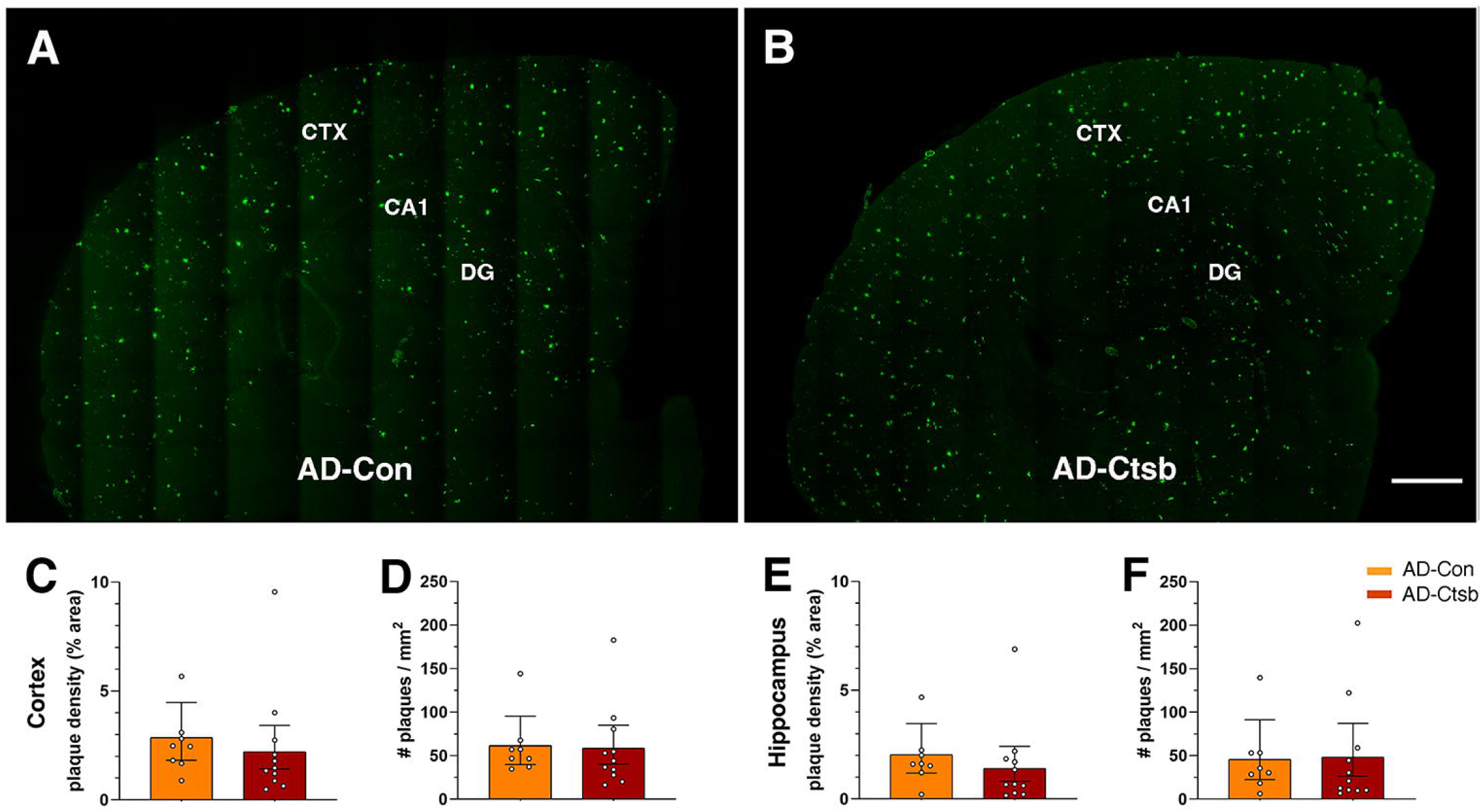
Ctsb treatment does not change amyloid plaque deposition. **(A,B)** Representative photomicrographs of ThioS staining (green) in sections derived from **(A)** AD-Con and **(B)** AD-Ctsb brain tissue. Scale bar: 500 μm. **(C-F)** There was no difference between AD-Ctsb and AD-Con mice in ThioS^+^ amyloid plaque density or number in **(C,D)** cortex or **(E,F)** hippocampus. Data were analyzed by GLM and are presented as Estimated Marginal Means and their respective 95% CI. (N; AD-Con = 8, AD-Ctsb = 11), Cortex (CTX); Dentate gyrus (DG).

To quantify microglia and astrocyte density, sections derived from the WT and AD groups were stained for Iba1 and GFAP, respectively (Figure 5A-D). Our analysis revealed a genotype, but not a treatment effect on Iba1 density. There was an increase in Iba1 (Figure 5E; Table S21) in AD-Con and AD-Ctsb as compared to WT-Con and WT-Ctsb groups. There was no group difference in GFAP density (Figure 5F; Table S22). Thus, our results show that Ctsb expression in AD mouse skeletal muscle does not affect plaque density or microglial activation.

**Fig 5.**
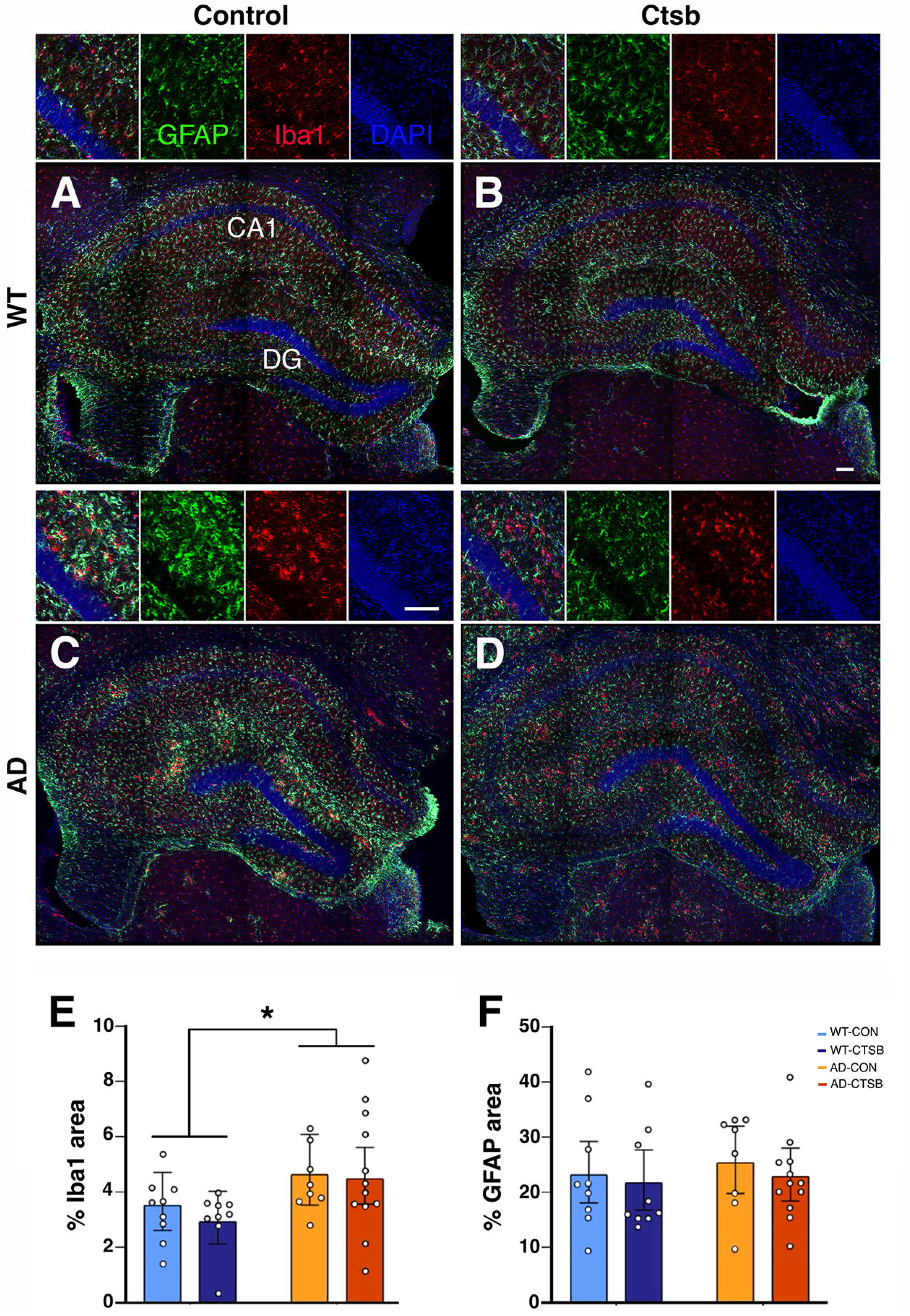
Ctsb treatment does not modify neuroinflammation. Representative photomicrographs of dorsal hippocampal sections subjected to GFAP (green), Iba1 (red) and DAPI (blue) labeling derived from (**A**) WT-Con, (**B**) WT-Ctsb, (**C**) AD-Con and (**D**) AD-Ctsb mice. **(E,F)** Astrocyte and microglia area density. Increased hippocampal density of **(E)** Iba1^+^ microglia but not of **(F)** GFAP^+^ astrocytes was observed in the AD groups. (N; WT-Con = 9, WT-Ctsb = 9, AD-Con = 8, AD-Ctsb = 12). Scale bars: 100 mm. *P<0.05 for Genotype effect (WT vs AD). Data were analyzed by GLM and are presented as Estimated Marginal Means and their respective 95% CI.

### Possible mechanisms mediating Ctsb treatment effects: proteomic analyses

To gain better insights into the mechanisms driving the effects of Ctsb treatment in AD on behavior and adult neurogenesis, proteomic analyses of hippocampus, muscle and plasma were performed (Figure 1B).

### Ctsb treatment enhanced transcription and translation in AD hippocampus

To investigate the potential effects of muscle-specific Ctsb treatment on the hippocampus, we conducted proteomic analysis on hippocampal tissue derived from Ctsb- and Con-treated AD and WT mice. Our workflow (Figure 1B) enabled the quantification of 8132 proteins, with 7509 proteins reliably quantified in more than three samples per experimental group (Figure S4B). Protein abundance ranged over approximately four orders of magnitude, encompassing highly abundant neuronal (e.g., App, Bdnf, Ckb, Thy1), cytoskeletal (e.g., Acta1.Actg2, Tuba1c) and glial (e.g., Gfap, and Apoe) proteins (Figure S4C). Multidimensional scaling showed no clear clustering pattern by genotype or treatment (Figure S4D).

To gain insight into the changes driven by AD pathogenesis, we first performed differential abundance analysis of average effect of AD. Only four proteins were significantly upregulated in AD, with a false discovery rate cut-off of 0.10 (Figure 6A): Amyloid precursor protein (App), vitronectin (Vtn), dedicator of cytokinesis protein 9 (Dock9) and complement C1q, subcomponent subunit C (C1qc) (Table S23). These proteins are well-known to be dysregulated in AD pathogenesis[44-47]. Our findings align well with the AD mouse model utilized in this study which overexpresses App, resulting in increased levels of APP and its peptide, amyloid beta-peptide 42, deposited in amyloid plaques[40]. Gene set enrichment analysis (GSEA) revealed that biological processes related to immune response, glycoprotein metabolic processes, amyloid fibril formation and glial cell development were significantly increased in AD mice (Figure 6B, Table S24). In contrast, DNA-templated transcription initiation was decreased (Figure 6B). GSEA based on cellular components showed that protein-lipid complex, lysosome and late endosome were significantly increased in AD, while synaptic membrane, potassium channel and transcription regulator complexes were decreased (Figure 6C). Some of the proteins associated with lysosome included the classical markers of AD, such as App, glial fibrillary acidic protein (Gfap), apolipoprotein E (Apoe) and nicastrin (Ncstn) (Figure 6D).

**Fig 6.**
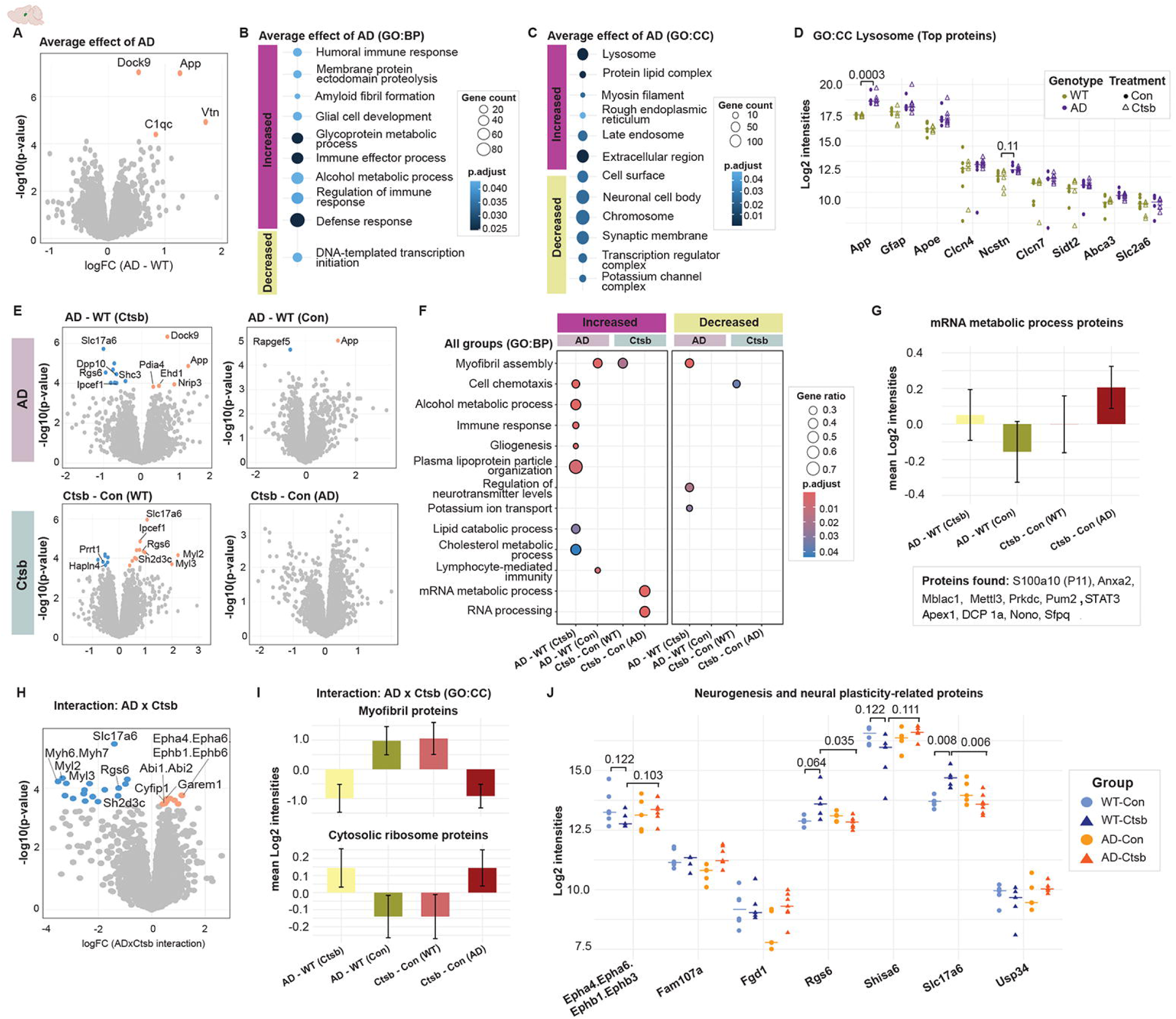
Ctsb treatment induces cytoskeletal reorganization in WT mice whereas transcription and RNA processing in AD mice. **(A)** Volcano plot of average effect of AD. Gene set enrichment analysis (GSEA) based on **(B)** biological processes and **(C)** cellular components. **(D)** Scatter plots of proteins associated with the lysosome ontology that has log fold changes > 0.50. **(E)** Volcano plots showing significant proteins in pairwise group comparisons. **(F)** GSEA of biological processes summarizing changes observed in all groups. **(G)** Mean abundance of proteins associated with mRNA metabolic process ontology. **(H)** Volcano plot of interaction between AD and Ctsb treatment. **(I)** Mean abundance of proteins associated with myofibril and cytosolic ribosome ontologies, which were found to be enriched by GSEA of interaction data, using cellular components. **(J)** Scatter plots of selected proteins from our data that play roles in neurogenesis and neural plasticity. Multiple hypothesis testing was performed by Benjamini-Hochberg method for each differential expression analysis and proteins with FDR < 0.10 were considered statistically significant. For GSEAs, a cut-off of FDR < 0.05 was used.

Next, we performed differential abundance analyses between each group (Table S23). The impact of AD pathogenesis was assessed by comparing AD-Con vs WT-Con [AD-WT(Con)], and AD-Ctsb vs WT-Ctsb [AD-WT(Ctsb)] groups, while WT-Ctsb vs WT-Con [Ctsb-Con(WT)] and AD-Ctsb vs AD-Con [Ctsb-Con(AD)] group comparisons were used to assess the effect of Ctsb treatment (Figure 6E-6F). Significantly regulated proteins for each group comparison, if present, are shown in Figure 6E (FDR < 0.10). GSEAs showed that most changes were specific to AD-Ctsb vs WT-Ctsb group, indicating a strong phenotype of the disease model, consistent with the behavioral findings where Ctsb treatment has opposite effects in WT and AD mice (Figure 6F, Table S24). Enrichment of ontologies such as increased immune response, gliogenesis and alcohol metabolic process in AD-Ctsb vs WT-Ctsb, recapitulated the average effect of AD as shown in Figure 6B. However, several processes showed opposing regulation; myofibril assembly increased in WT-Ctsb and AD-Con mice, but decreased in AD-Ctsb and WT-Con groups. In addition, cell chemotaxis was diminished in WT-Ctsb but elevated in AD-Ctsb mice. Moreover, Ctsb treatment increased mRNA metabolic process and RNA processing in AD-Ctsb vs AD-Con mice (Figure 6F). Further investigations into the mRNA metabolic process revealed that abundance of proteins associated with this ontology increased, especially in AD-Ctsb vs AD-Con group (Figure 6G). Some of the proteins involved here were S100a10 (also known as P11) and annexin A2 (Anxa2), which form a complex that interacts with various ion channels and receptors. Alterations in S100a10 levels are observed in cancer, psychiatric and neurodegenerative disorders[48]. In fact, we previously showed that S100a10 is required for the beneficial effects of Ctsb on neuronal differentiation and migration[24].

Lastly, differential abundance analysis exploring the interaction between AD and Ctsb showed 17 decreased and eight increased proteins (FDR < 0.10) (Figure 6H, Table S23). GSEA of interaction data showed that cellular components such as myofibril and actin cytoskeleton were decreased while the component cytosolic ribosome was increased (Table S24). Mean abundance of proteins associated with these ontologies clearly showed that AD-Con and WT-Ctsb groups had similar overall protein abundance as compared to WT-Con (Figure 6I). Ctsb treatment in AD mice showed a decreased tendency in overall abundance of proteins in myofibril whereas an increased tendency of cytosolic ribosomal proteins. It is noteworthy that, following Ctsb treatment, proteins typically found in muscle, such as myosin light chain (Myl2 and Myl3), were decreased in hippocampus of AD mice, but upregulated in the hippocampus of WT mice (Figure 6E and 6I). This observation aligns with earlier observations where abundance of muscle proteins increased in brain during aging and in pathological conditions. For instance, in human AD brain, a muscle protein, dysferlin, is associated with amyloid plaques[49], while Myl9 is upregulated in the aging human hippocampus[50]. Interestingly, several proteins that are involved in neurogenesis and neural plasticity were regulated similarly in AD-Ctsb and WT-Con, as compared to WT-Ctsb and AD-Con groups (Figure 6J). These findings support that Ctsb treatment normalises brain function in AD mice.

### Ctsb treatment decreased WT muscle mitochondrial processes, whereas it increased translation in AD muscle

To characterize the effect of muscle-specific Ctsb treatment, we performed muscle proteomics and quantified 2998 proteins after stringent filtering (Figure S4E). Multidimensional scaling plots, which did not exhibit any indications of clustering patterns, are shown in Figure S4F and S4G.

Differential abundance analyses between groups did not reveal significant changes in proteins with the cut-off of FDR < 0.10 (Table S25). Ctsb abundance did not differ between groups eight months post-treatment, suggesting return to- or maintenance of physiological levels (Figure S4). Nevertheless, GSEA of the average effect of AD pathogenesis showed an increase in Wnt signaling pathway, with APP being the most increased protein (Figure 7A, Table S26). The Wnt signaling pathway, crucial for cell differentiation and muscle regeneration in adult muscle[51], is also known to regulate APP processing and is dysregulated in the brain during AD[52]. Elevated APP levels within muscle tissue have been suggested to play a role in brain AD pathology in APP/PS1 mice[53]. GSEA of the average effect of Ctsb treatment suggested an increase in Erk1-Erk2 cascade, translation at presynapse and defense response and a reduction in cellular respiration, mitochondrial translation and transport (Figure 7B). Similarly, cellular component ontologies showed a decrease in mitochondrial terms (Figure S4L).

**Fig 7.**
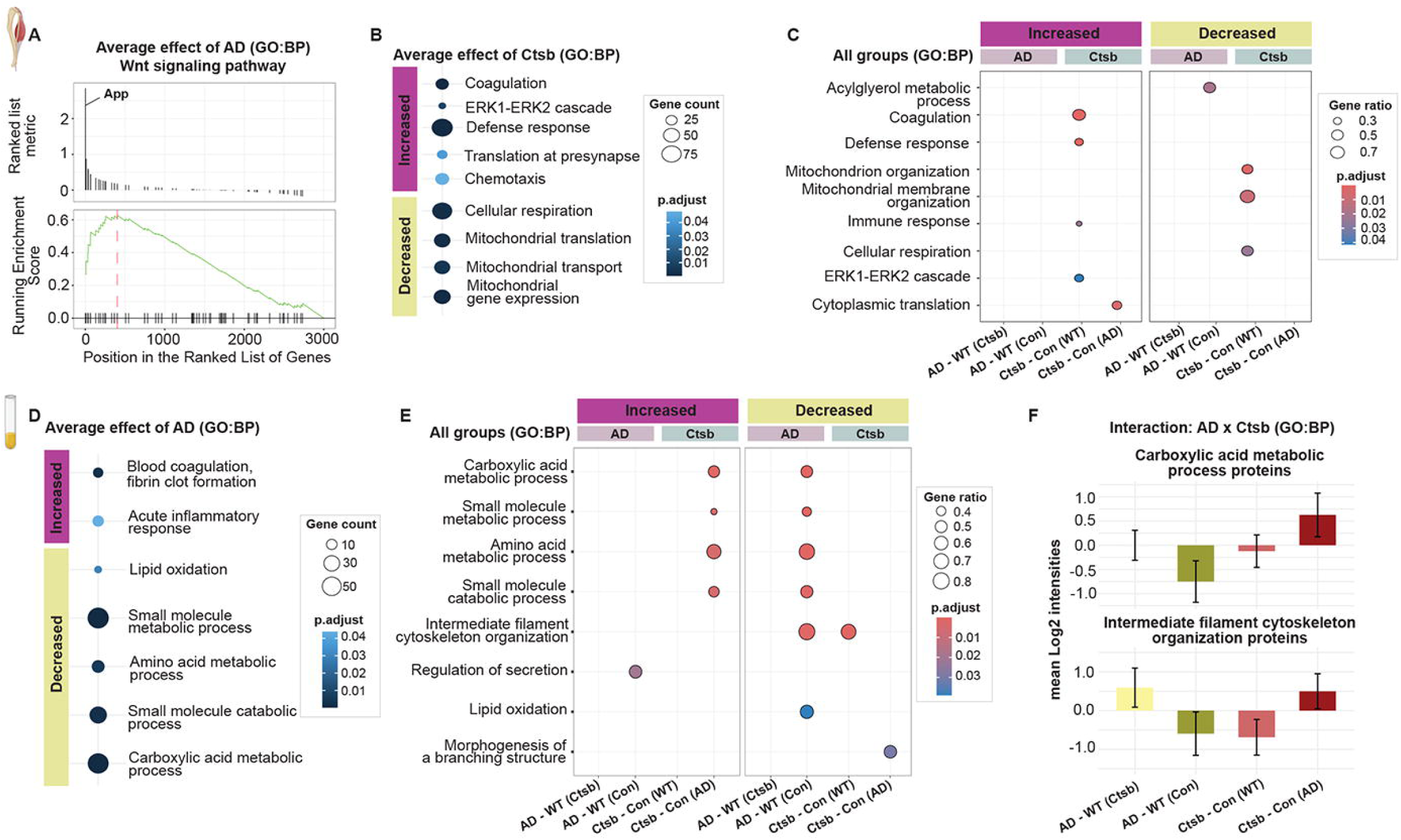
Effects of Ctsb treatment on muscle and plasma proteome. Muscle proteome: **(A)** GSEA plot of Wnt signaling pathway, as one of the most affected gene set in AD. GSEA of average effect of Ctsb treatment based on **(B)** biological processes. **(C)** GSEA using biological processes summarizing changes observed in each group. Plasma proteome: GSEA of **(D)** biological processes summarizing changes observed in the average effect of AD. **(E)** GSEA of biological processes summarizing changes observed in all groups. **(F)** Mean abundance of proteins associated with metabolic and cytoskeletal ontologies were displayed from GSEA of interaction data. **(C,E,F)** Data for the pairwise comparisons are annotated as follows: AD-WT(Ctsb), *AD-Ctsb vs WT-Ctsb*; AD-WT(Con), *AD-Con vs WT-Con*; Ctsb-Con(WT), *WT-Ctsb vs WT-Con*; Ctsb-Con(AD), *AD-Ctsb vs AD-Con*.

GSEA, performed in each pairwise group comparisons, suggested that many enriched ontologies were prominent in Ctsb-treated WT mice (Figure 7C). Increased defense response, coagulation and Erk1-Erk2 cascade as well as reduced mitochondrial organization, transport and matrix were observed in WT-Ctsb mice. In contrast, increased translation was unique to AD-Ctsb mice. Similar observations were found by cellular components (Figure S4M). These findings underscore the significant interplay between muscle-derived proteins and brain pathology, suggesting that alteration in muscle proteome may play a crucial role in the progression of Alzheimer’s disease.

### Metabolic processes in AD plasma are increased by Ctsb treatment

To elucidate the systemic effects of Ctsb treatment in the context of Alzheimer’s disease, we conducted a comprehensive plasma proteome analysis. Our workflow (Figure 1B) led to identification of 1132 proteins of which 713 were quantified in more than three samples per group, indicating that our protocol resulted in a good coverage (Figure S4H). The classical high abundant plasma proteins (e.g. albumin, serpinas, apolipoproteins) were highlighted in the dynamic range curve, which covered four orders of magnitude (Figure S4I). Sample distributions did not show genotype- or treatment-specific clustering (Figure S5J).

Sodium/potassium-transporting ATPase subunit gamma (Fxyd2) was the only altered protein (FDR < 0.10) by differential abundance analysis exploring the average effect of AD (Table S27). This protein is likely to be secreted from immune cells because the Fxyd2 mRNA transcript is primarily expressed by memory CD8 T-cells and natural killer cells (https://v19.proteinatlas.org/ENSG00000137731-FXYD2/blood). GSEA based on biological processes suggested that metabolic and catabolic processes were decreased in AD (Figure 7D, Table S28). In contrast, blood coagulation and acute inflammatory response terms were increased, which are known to be altered in AD pathogenesis[54]. Interestingly, proteins associated with mitochondrial terms were reduced in AD (Figure S4N). However, group-based changes showed that overall protein abundance of mitochondrion had an increased tendency upon *Ctsb* expression in AD group (Figure S4L). GSEA of each pairwise group comparison showed that gene sets related to metabolic and catabolic processes were positively enriched in AD-Ctsb vs AD-Con group whereas negatively enriched in AD-Con vs WT-Con group (Figure 7E, Table S28). Thus, the treatment brought proteomic profile of the AD mice closer to that of the WT control group. Lastly, differential abundance analysis of interaction between treatment and genotype was performed. GSEA led to enhanced enrichment of metabolic processes and intermediate filament-based process (Table S28). Mean abundance of proteins associated with these ontologies clearly showed that AD-Con vs WT-Con and WT-Ctsb vs WT-Con groups had similar overall protein abundance (Figure 7F). These results highlight that Ctsb treatment not only influences muscle and hippocampal proteome but also enacts systemic metabolic changes that may contribute to ameliorating Alzheimer’s disease symptoms.

## DISCUSSION

Increasing evidence for a role of peripheral organs in brain function has opened novel opportunities to prevent and treat neurodegenerative diseases. We hypothesized that long-term expression of *Ctsb* in muscle, starting from around the onset of disease, may ameliorate AD-related pathologies in APP/PS1 mice. Here we show that this treatment substantially improves memory, motor behavior, and adult hippocampal neurogenesis in AD, but not in WT mice. Proteome analysis of hippocampus, muscle and plasma revealed specific remodeling associated with AD and Ctsb treatment. These changes may contribute to increased adult neurogenesis and thereby improve memory function in Ctsb treated AD mice.

Middle-aged APP/PS1 mice display deficits in learning, memory and motor function[55], behaviors mediated at least in part by the hippocampus[56, 57]. Impaired rotarod performance in AD was observed, consistent with earlier studies[58], and was reversed by muscle *Ctsb* expression. In addition, although AD mice acquired the water maze task slower than WT mice, retention of spatial memory was substantially improved by treatment, comparable to exercise interventions in middle-aged mice[59]. On the other hand, WT mice treated with Ctsb performed at chance levels in the probe trials suggesting a detrimental effect of treatment on retention of spatial memory. Similarly, in the fear memory task, the WT-Ctsb group did not condition and displayed reduced cued fear, suggesting disruption of the relevant hippocampal, amygdala and prefrontal cortex circuitry[60, 61]. While reduced freezing behavior could be attributed to increased general activity, as observed in the activity box, the impaired retention of spatial memory indicates that poor fear conditioning performance is likely due to cognitive deficits. The other groups displayed tone-shock conditioning and cued fear, as observed in other aging[62] and AD mouse[63] studies. Contextual conditioned fear reaction improved in treated AD mice as it was comparable to wildtype controls. In the AD control group contextual freezing may reflect over-generalization[64]. Overall, Ctsb expression in AD mice resulted in spatial and fear memory behavior that is largely similar to that of the wildtype control group.

Improved memory function in AD mice treated with Ctsb may be due in part to increased adult hippocampal neurogenesis, a highly regulatable process relevant to cognition and mood. Adult neurogenesis declines in normal aging, and more substantially in AD mice, and has been suggested to play a role in human disease[65]. Reduced cell genesis is generally observed by eight to ten months of age in APP/PS1 mice[66]. In our study, doublecortin cell numbers at twelve months of age were decreased in the AD control group. Ctsb treatment prevented this deficit in AD mice, possibly by modulation of gene expression and protein activity within the neurogenic niche[67, 68]. Our proteomic analyses showed that in hippocampi derived from treated AD mice RNA metabolism is increased. Specifically, the transcription regulator complex was modified in a manner that is opposite to AD controls. For instance, hippocampal P11 (S100A10), a protein that regulates neurogenic effects of Ctsb[24], and its binding partner Anxa2[69] increased only in the Ctsb-treated AD group. Signal transducer and activator of transcription 3 (STAT3), an epigenetic regulator of neurogenic processes[67, 70], and Nono, a protein important for cell cycle and synaptic function[71] had similar increased tendency. It is also noteworthy that Sfpq a RNA binding protein with roles in transcriptional elongation, mRNA processing and DNA repair that is downregulated in human AD patient brains[72], was elevated, albeit not significantly, by Ctsb treatment in AD mice. Furthermore, several proteins involved in neurogenesis and neural plasticity were regulated similarly in wildtype controls and Ctsb treated AD mice, including Slc17a6[73], Rgs6[74, 75], Epha4.Epha6.Ephb1.Ephb3[76] and Shisa6[77], potentially supporting improved memory function.

Differences between AD and WT in muscle proteome in response to Ctsb treatment may also contribute to behavioral and neurogenic outcomes. In WT muscle Ctsb caused a decrease in mitochondrial and increased coagulation processes, which may be linked to deficient cognition [6]. With aging and neurodegenerative disease skeletal muscle mass is reduced[4, 6]. Strength training reportedly improves cognitive function in the APP/PS1 mouse model[78]. Our analysis shows increased cytoplasmic translation and cytosolic ribosome with Ctsb treatment in AD mice. Improved rotarod performance may be linked in part to these changes, as these processes are important for muscle growth and maintenance[79]. Furthermore, in plasma mitochondrial abundance had an increased tendency, whereas plasma inflammatory responses[80] were reduced in AD, suggesting that Ctsb has a disease-modulating role.

The current approaches to AD treatment are focused on diminishing neuroinflammation and reducing amyloid plaques[2, 3]. While these are important targets, our study shows that even in the absence of changes in plaque pathology, APP levels and neuroinflammation, muscle Ctsb treatment promotes adult hippocampal neurogenesis and memory function. Moreover, our proteomics analyses across tissues and plasma provides insight into the potential underlying mechanisms, further supporting the concept that there are novel and thus far sparsely explored signaling pathways that could be harnessed to treat the disease. However, our study has several limitations and there are remaining questions that need to be addressed. The treatment benefits AD mice but is detrimental in wildtype mice. It remains to be determined why, however, the proteomics analyses show increased coagulation and decreased mitochondrial function in muscle, and reduced cell chemotaxis and increased cytoskeleton pathways in hippocampus, which together may reduce neural plasticity. Some limitations include use of male mice only and reduced power due to limited sample size, however, given the promise of the results for AD intervention, further research is warranted. Furthermore, it will be important to investigate if the present findings can be extended to other AD mouse models and whether the shorter or longer time-course of administration of the vector impacts outcome. Finally, it is unknown if targeting organs other than muscle would have similar or different outcomes for the measured parameters.

Altogether, muscle Ctsb treatment in AD mice benefits adult neurogenesis and memory function, despite the presence of plaque pathology and neuroinflammation. Proteomic analysis across tissues and plasma revealed peripheral and central changes that may underlie the disease-modifying outcomes. Muscle Ctsb has potential as a therapeutic approach for neurodegeneration.

## MATERIALS AND METHODS

### Mice and Housing Environment

B6.Cg-Tg(APPswe,PSEN1dE9)85Dbo/Mmjax (APP/PS1) double transgenic hemizygous male mice on the C57BL/6J background, and female mice without the APP/PS1 allele (referred to here as wild-type; WT) were purchased from Jackson Laboratories (MMRRC Strain #034829-JAX). Mice were group housed in Individually Ventilated Cages (Tecniplast, Emerald line) containing bedding and nesting material (nestlets), and acclimated to the vivarium environment prior to breeding. Cages were maintained in a temperature-controlled environment (21 ± 2°C) and under a 12h/12h light-dark cycle (lights were switched off at 7:30 PM), with water and food available *ad-libitum*.

All animal-use procedures were conducted after approval by Florida Atlantic University’s Institutional Animal Care and Use Committee and they were conducted in accordance with the National Institutes of Health Guidelines for the care and use of Laboratory Animals.

### Experimental Procedures

#### Breeding strategy and allocation to the groups

WT female and APP/PS1 hemizygous male mice (6-8 months old) were used as breeding pairs to produce APP/PS1 hemizygous mice (which were utilized as the AD model) and their WT littermates (serving as genotype controls). A 1:1 female-to-male ratio was employed as an in-house breeding strategy. Litters were with their respective dams from the day of birth (postnatal day zero - PND0) until weaning at PND20-21. The weaning procedure involved ear-tagging the pups, collecting tissue (tip of the tail) for genotyping (Transnetyx), and group housing the mice (3-4 mice per cage) according to sex. No more than two mice per genotype/litter were assigned to the same treatment group. Four-month-old male mice, derived from 2 cohorts of mice, were allocated to one of the following groups for either Ctsb or Control treatment: WT-Con (N=10 (6+4)), WT-Ctsb (N=9 (6+3)), AD-Con (N=8 (5+3)), and AD-Ctsb (N=14 (10+4)).

#### Outline of experiments

At four months of age, AD mice and their WT littermates, were allocated to either Ctsb or Control treatment. Ctsb was over-expressed in muscle tissue by injecting an AAV vector into the tail vein, to express the mouse *Ctsb* gene driven by the muscle creatine kinase promoter (MCK). Upon five months of age mice were injected with bromodeoxyuridine to label dividing cells. Mice were left undisturbed until the onset of behavioral testing at 10 months of age (as detailed below). Upon conclusion of behavioral testing, mice were 11.5-months old. At 12 months of age the animals were deeply anesthetized for blood and tissue collection.

##### AAV Vector

Four-month-old mice received tail vein injections of pAAV9-tMCK-mCTSB-IRES-eGFP or control vector pAAV9-tMCK-eGFP-WPRE (VB5037) (Vector BioSystems). For tail vein injections, mice were lightly anesthetized within an isoflurane chamber with isoflurane (Abbott) and subsequently placed onto a tail illuminator restrainer (Braintree Scientific). Once the ventral tail vein was dilated by the heat of the illuminator, 50 μL of vector (10^11 vg/ml) was slowly injected into the blood vessel. When mice were fully recovered from anesthesia (∼10 min) they were returned to their home cage.

##### Bromodeoxyuridine (BrdU)

One month after vector treatment, 5-month-old mice received intraperiotoneal injections of BrdU (50 mg/kg; 5 ml/kg) for 10 consecutive days. BrdU (Sigma, B5002-1G) was prepared fresh daily by dissolving the powder in 0.9% saline at a 10 mg/mL concentration, followed by 30 min incubation in a 37 °C water bath, filtration through a sterile syringe at 0.2 µm, and was protected from the light prior to injection.

### Behavioral tests

Upon 10 months of age mice were subjected to a battery of behavioral tests. Mice were tested in the activity box (open field) test, rotarod, Morris water maze, and fear conditioning. All tests were performed during the light period of the light-dark cycle.

#### Activity Box (Open field)

Tests were carried out in empty plexiglass arenas (height 20.3 cm, width 27.2 cm, depth 27.3 cm) containing two 16-photo beam infrared (IR) arrays on the X and Y axes. Eight arenas were used simultaneously, each one located inside a sound-attenuated chamber equipped with two ceiling white lights (SKU ENV-221CL, Version 4.0, Med Associates, St. Albans, VT, USA). Mice were acclimated to the testing room for at least 45 min before starting the test. Procedure consisted of placing the mice in the center of the arena and allowing them to move freely for 60-minutes. Spontaneous locomotor activity (ambulatory distance) was recorded by Activity Monitor software (Med Associates).

#### Rotarod

Motor function was evaluated with an accelerating rotarod for mice (Med Associates, St. Albans, VT). The apparatus consisted of a rotating drum (3.2 cm diameter) separated by white plexiglass matte walls, forming five 8 cm wide stations that allows for simultaneous testing of five animals. Mice were acclimated to the testing room for at least 30 min before starting the test. Procedure consisted of placing the mice on the rotating cylinder, which was turned on at a constant low speed (4 rpm). Once all five animals were positioned on the rotating cylinder, the speed was shifted from the constant 4 rpm to an accelerating speed by adding 0.12 rpm/second, changing from 4 to 40 rpm over five minutes. The latency to fall was recorded by Activity Monitor software (Med Associates). If an animal did not fall during the five minutes of testing, a 300s latency to fall was scored for that animal.

#### Morris water maze

Spatial learning and retrieval were evaluated using the Morris water maze paradigm[42]. The apparatus consisted of a pool (1.83m diameter) filled with water (24-26 °C), made opaque with white nontoxic paint (Tempura, Crayola), and contained a platform (20 cm x 20cm) that was hidden 1 cm below the surface of the water. The pool was surrounded by a black curtain, 60 cm from the edge of the pool to which visual cues were attached. Indirect illumination of both the pool and cues (65 lux at the center of the maze) was achieved by positioning 4 white lights equidistantly around the pool and facing the ceiling. The pool was virtually separated into 4 quadrants, called northeast (NE), northwest (NW), southeast (SE) and southwest (SW). Mice were acclimated to the testing room daily for at least 30 min before starting the test. Acquisition sessions consisted of four 60-s trials with a 15-s inter-trial interval, for 20 days, in which the hidden platform was located in the NE quadrant of the pool. Mice were placed at a different starting point for each trial. If the mouse found the platform, the trial was immediately terminated. Mice were placed on the platform if they did not find it within the allocated time. In either case, mice were allowed to explore the visual cues for 15-s while on the platform. Spatial memory retention was evaluated throughout two 60-s probe trials (performed 4 h and 24 h after completion of the last acquisition session) in which the platform was removed. For both probe trials, mice started in the SW quadrant. Data was collected by an automated video tracking system (Ethovision XT 17.5).

#### Fear conditioning paradigm

Pavlovian conditioning-based approach was used to evaluate fear memory. Mice were tested within a four-chamber system utilizing a near-infrared video conditioning system (MED Associates Inc., Fairfax, VT, USA) that allows for testing of four animals simultaneously. Each chamber consists of stainless-steel walls with one transparent plexiglass door and a floor of parallel stainless-steel rods connected to a shock generator. The chamber has an overhead white light (SKU ENV-229M) to illuminate the inside (3 to 8 lux), and a speaker which is used to deliver a tone, to serve as the conditioned stimulus – CS. Each of these chambers was located within a larger noise-attenuating chamber (height 31.75 cm x width 71.12 cm x depth 59.69 cm), including a ventilation fan delivering background noise. A near-infrared camera-tracking system (MED Associates, Georgia, VT, USA) was used to automate measurement of freezing behavior (Motion Threshold (au) = 20; Detection Method = Linear; Min Freeze Duration (f) = 18). The paradigm consisted of four phases: (i) Habituation, (ii) Conditioning, (iii) Tone-cued, and (iv) Contextual phase. Mice were acclimated to the testing room for at least 1 h before test onset. Habituation (Day 1) consisted of a single 330-s session in which mice were placed within the chamber and allowed to freely explore the environment. A 1% liquinox solution was in the tray located underneath the stainless-steel rods to serve as an odor cue and to facilitate contextual perception and discrimination. Freezing behavior was measured for the entire 330-s. Mice were immediately returned to their home cages at the end of the session. The inner surfaces of the chamber and the tray were cleaned thoroughly with 70% isopropyl alcohol. Conditioning phase (Day 2) was similar to the Habituation session with the exception that a CS, tone, paired with an Unconditioned Stimulus (US), foot shock, was presented 60 seconds after the beginning of the trial. CS consisted of a tone (5000 Hz, 90 dB tone), for 30 s, that ended with the presentation of the US, a 0.5s foot shock (0.5mA) delivered 0.5s before the end of the CS (TS1). Thereafter, the CS-US was repeated two more times (TS2 and TS3, respectively) with a 90-s inter-stimulus interval. Freezing behavior was measured during the same 30s in which the tone was on. CFR, the response evoked by the CS-US association, was considered to be formed if freezing behavior was higher during the third than the first CS-US presentation. Tone-cued phase (Day 3) was performed to evaluate long-term evocation of Pavlovian conditioning. It consisted of a single session similar to the Conditioning phase, except that no US was delivered. In addition, to avoid contextual (visual, tactile and olfactory) influence on the CS-US association, the original environment was modified by adding a blue semitransparent plastic film on top of the plexiglass chamber, an arch-shaped white plastic panel within the chamber and a white plastic panel at the base of the chamber as well as the replacement of the 1% liquinox solution by an 1% acetic acid solution. Three Contextual sessions (Day 3 - one hour after Tone-cued phase – Day 5 and Day 7; named C1, C2 and C3, respectively) were performed to evaluate long-term evocation of contextual CFR (i.e., whether freezing behavior was higher during the tests than during Day 1). These were identical to the Habituation session.

### Euthanasia and Tissue collection

Upon completion of the behavioral experiments, mice (11.5 months-old) were deeply anesthetized for terminal blood and tissue collection. Approximately 300 µL of trunk blood was collected from mice into EDTA-coated tubes and directly placed on ice. Within 15 minutes these samples were spun at (2,000 RCF) for 15 minutes within a temperature controlled (4 °C) centrifuge. Supernatant was collected and 2-3 aliquots (50-75 µL) were stored at -80 °C until further use. Following transcardiac perfusion with 0.9% saline (room temperature), liver, gastrocnemius, soleus, biceps and heart tissue were dissected, immediately flash-frozen in liquid nitrogen, placed on dry ice, and stored at -80 °C. Half of the brain was dissected in a petri dish over cold ice to collect the prefrontal cortex and hippocampus. For the present study, hippocampus, gastrocnemius and plasma were utilized for proteomic analyses. The other hemisphere was placed in ice cold 4% paraformaldehyde (PFA), post-fixed for 72 hours, and subsequentially equilibrated in 30% sucrose until sectioning.

### Histology

Sequential coronal sections (40 μm) were taken using a freezing microtome (HM450, ThermoFisher Scientific, Wyman St. Waltham, MA, USA) throughout the whole rostrocaudal extent of one hemisphere of the brain, and stored in 96-well plates in a phosphate-buffered glycerol anti-freezing solution at -20 °C, as described[81].

#### Immunohistochemistry

For both DCX and BrdU staining, sections were subjected to the following: (i) quenching endogenous peroxidase activity, (ii) antigen retrieval, (iii) immunolabeling, (iv) detection and dehydration.

#### DCX staining

A one-in-twelve series (480 μm apart) of free-floating sections was rinsed in 0.1M Tris-Buffered Saline (TBS, 3 x 5-min). (i) sections were incubated for 30 min (at room temperature (RT)) in 0.6% H_2_O_2_ in TBS to quench endogenous peroxidase activity. Five-min rinses in TBS were performed until no bubbles were detected. (ii) Antigen retrieval: incubation in 1mM citrate buffer (pH 6.0) at 95-98 °C for 20 min. (iii) Sections were rinsed 3 x 5-min in TBS followed by 1h (at RT) in blocking solution (3% donkey serum and 0.1% Triton x-100 in TBS (TBS^++^)), and overnight incubation at 4 °C in TBS^++^ containing the primary antibody (mouse monoclonal anti-DCX, 1:1000, Santa Cruz Biotechnology Cat# sc-271390, RRID:AB_10610966). Next, immunolabeling steps were repeated, albeit sections were incubated for 2h at RT with the secondary antibody (Biotin-SP goat anti-mouse, 1:500, Jackson ImmunoResearch Labs Cat# 115-065-166, RRID:AB_2338569). (iv) The detection steps: TBS (rinses 3x 5 min) followed by 2h incubation (at RT) in avidin-biotin-peroxidase complex (VECTASTAIN® Elite® ABC-HRP Kit - Vector Laboratories), TBS (3x 5-min), a 7 min incubation in diaminobenzidine (DAB) chromogenic substrate and urea hydrogen peroxide (Sigma-Aldrich # D4418), and TBS (5x 5-min) rinses. Sections were mounted onto gelatin subbed slides, dehydrated using increasing alcohol concentrations and CitriSolv (Fisher Scientific 04-355-121), and cover-slipped with DPX mounting media (Sigma-Aldrich # 06522).

#### BrdU staining

A one-in-six series (240 μm apart) of free-floating sections were subjected to (i) quenching of endogenous peroxidase activity (as described for DCX staining). (ii) Antigen retrieval consisted of incubating the sections in 50% formamide diluted in 2x SSC buffer (30mM sodium citrate in 0.3M NaCl) at 65 °C for 2 h, followed by 5 min incubation in 2X SSC buffer at RT, 60 min in 2N HCl at 37 °C, and 10 min incubation in 0.1M borate buffer (pH 8.5) at RT. (iii) Immunolabeling and (iv) detection and dehydration steps were identical to DCX staining, except for the primary (rat monoclonal anti-BrdU, 1:500, Abcam Cat# ab6326, RRID:AB_305426) and secondary antibodies (Biotin-SP polyclonal donkey anti-rat, 1:500, Jackson ImmunoResearch Labs Cat# 712-065-153, RRID:AB_2315779).

DCX^+^ cells and BrdU^+^ cells were quantified from the rostral to caudal dentate gyrus. Detailed information is presented in Supplementary Material (Supplementary Information S3). In order to minimize detection bias, a computer-based random order generator was used to predefine the order in which each slide was analyzed (slide IDs were coded) by an experimenter who was blinded to the groups. The average number of positive cells per section was calculated.

#### Thioflavin-S (ThioS) Staining

ThioS working solution (Sigma-Aldrich: T1892) was prepared at a final 0.01% concentration diluted in 50% ethanol. A one-in-twelve series of free-floating sections (40 μm) was rinsed in TBS (3 x 5-min) followed by 30 min incubation in 0.01% ThioS solution, 5 min incubation in 70% ethanol, and rinses in dH_2_O (3 x 5 min). Sections were mounted on gelatin subbed slides, and coverslipped with DABCO-PVA.

Sections were systematically imaged on an epifluorescence microscope (Olympus BX51) and quantified. For each section, one image was taken of the following brain areas: the dentate gyrus (DG) (granule cell, polymorphic and molecular layer), CA1 (pyramidal cell layer, stratum oriens, radiatum, and lacunosum moleculare), auditory/somatosensory cortex (primary/secondary auditory/somatosensory cortex), perirhinal/entorhinal cortex (perirhinal and/or dorsolateral entorhinal cortex) and cingulate cortex (retrosplenial dysgranular cortex and retrosplenial granular cortex). The “interactive learning and segmentation toolkit” [82] was used to conduct unbiased image segmentation. Briefly, after extensive training of Ilastik to identify pixels as either positive- or negative-ThioS labeling (background), a pixel-based classification was applied to all images. A ThioS^+^ plaque was defined as any green fluorescent labeling above the background (using an image derived from WT tissue as ThioS-negative comparison – background control). Binary output images of ThioS^+^ labeling were exported through the “simple segmentation” feature. The “Measure” feature on ImageJ software (Image J2: Version 2.14.0/1.54f) was used for collecting data from the simple segmentation images. The estimation of ThioS^+^ plaque density (% area covered/0.32 mm^2^) was calculated either for each individual region and combined for hippocampus and cortex. ThioS^+^ plaques were also counted for each 0.32 mm^2^ area using the multi-point tool on ImageJ software (version 1.54g). A ThioS^+^ plaque was defined as any green fluorescent labeling above the background (using an image derived from WT tissue as ThioS-negative comparison – background control). For the cases in which two or more green fluorescent areas were connected to each other (forming a single “ramified” area), this was considered a single ThioS^+^ plaque. Finally, the estimation of the number of ThioS^+^ plaques/mm^2^ was calculated for each individual region and pooled for hippocampus and cortex. A computer-based random order generator used to predefine the order in which each slide was imaged and analyzed (slide IDs were coded) by experimenters who were blinded to the groups during the whole procedure.

#### GFAP and Iba1 immunofluorescence

A one-in-twelve series of free-floating sections (40 μm) was rinsed in TBS (3 x 5-min) and incubated for 1h in TBS^++^ (at RT). Thereafter, sections were incubated in TBS^++^ containing a cocktail of primary antibodies for 72h at 4 °C (rabbit polyclonal anti-GFAP, 1:1000, Dako Cat# Z0334, RRID:AB_10013382; goat polyclonal anti-Iba1, 1:500, Abcam Cat# ab5076, RRID:AB_2224402). After rinses with TBS (3x) the sections were incubated (at RT) in TBS^++^ containing their respective secondary antibodies for 2h (Alexa Fluor 488 donkey anti-rabbit, 1:500, Jackson ImmunoResearch Cat# 711-545-152, RRID: AB_2313584; Cy3 donkey anti-goat, 1:500, Jackson ImmunoResearch Cat# 705-165-147, RRID: AB_2307351). Sections were rinsed with TBS (3 x 5 min), incubated with DAPI solution (0.5 µg/ml; Molecular Probes: MP01306) for 5 minutes, rinsed (3 x 5 min), mounted onto gelatin-coated slides and cover-slipped with DABCO-PVA.

To quantify astrocytes (GFAP^+^) and microglia (Iba1^+^) area, tile scan images covering the entire hippocampal formation were acquired on a confocal microscope (Nikon, A1R). Similar to the procedure described in ThioS analysis, Ilastik (version 1.4.0.post1-gpu)[82] was used to conduct unbiased image segmentation on “max projection” confocal output images. Briefly, after extensive training of Ilastik to identify pixels as either positive or negative (background) immunofluorescent labeling, a pixel-based classification was applied to all images. To avoid miss-estimation of either GFAP^+^ or Iba1^+^ immunolabeling due to superimposition of pixels, independent training was conducted for each channel. Binary output images of either GFAP^+^ or Iba1^+^ immunolabeling were exported through the “simple segmentation” feature. The density of GFAP^+^ and Iba1^+^ labeling in the hippocampus was estimated by systematically sampling two 0.15 mm^2^ Region of Interests (ROI) per binary image, one in the DG (containing the polymorph layer, the granule cell layer and the molecular layer) and one in the CA1 subfield (containing the pyramidal cell layer, the stratum oriens, the stratum radiatum, and the stratum lacunosum moleculare). Unbiased ROI positioning was conducted using the DAPI channel as reference in ImageJ software (Image J2: Version 2.14.0/1.54f) and applied to their respective simple segmentation images for data collection utilizing the “Measure” feature. Finally, estimation of the density of GFAP^+^ and Iba1^+^ labeling (% area covered/0.15 mm^2^) was calculated either for each individual region or for all of them pooled together. Both performance and detection biases were minimized by using this approach combined with experimenters who were blinded to the groups to conduct the entire procedure.

### Proteomics

#### Sample preparation for proteomics analysis

Hippocampal and gastrocnemius muscle tissues were powdered and lysed in 4% Sodium Dodecyl Sulfate (SDS) buffer (100 mM Tris, pH 8.5) using Ultra Turrax blender (IKA). Lysates were then boiled at 95 ^0^C for 10 min, followed by 10 min sonication in ultrasonic bath sonicator. After centrifugation at 16000 g for 10 min, supernatants were transferred into a new tube for determination of protein concentration using DC protein assay (Thermo Fisher Scientific). Subsequently, 20 μg of proteins were reduced by the addition of 100 mM of dithiothreitol and alkylated by 40 mM of chloroacetamide. After 45 minutes of incubation at the room temperature, proteins were digested using PAC protocol on KingFischer Flex robot [83]. In brief, a 1:4 protein to bead ratio was added to the sample lysate and protein aggregation was induced by dispensing a final concentration of 70% acetonitrile (ACN), followed by a 10-min incubation without agitation. Then, the sample plates were placed in a magnetic stand (Dynamag Thermo) and beads with protein aggregates were first washed twice with 100% ACN and then twice with 70% ethanol. After discarding the last wash, beads were resuspended in 100 μl of digestion buffer (50 mM Tris-HCl, pH 8.5) containing 1:100 enzyme to protein ratio of trypsin and 1:500 enzyme to protein ratio of LysC (Wako) and enzymatic digestion was done at 37 °C overnight. The next day, digestion was quenched by addition of a final concentration of 1% trifluoroacetic acid (TFA) in isopropanol. Peptides were then cleaned for salts and remaining lipids using styrenedivinylbenzene–reverse phase sulfonate (SDB-RPS, details) stage-tips and eluted in 60 μl of 1% ammonia and 80% ACN. Using speedvac, peptides were dried completely and resuspended in 10 μl of 0.1% TFA and 5% ACN. Peptide concentration was determined by a NanoDrop spectrophotometer (Thermo Fisher Scientific) and a total of 200 ng of peptides were loaded on Evotip C18 trap columns (Evosep Biosystems) according to the manufacturer’s instructions.

For the plasma proteomics, 1 μl of plasma was directly lysed in 100 μl of 100 mM Tris buffer (pH 8.5). Lysates were then boiled at 95 ^0^C for 10 min, followed by 10 min sonication in ultrasonic bath sonicator. After a quick spin down, enzyme mix (containing 1:100 enzyme to protein ratio of trypsin and 1:500 enzyme to protein ratio of LysC) were added and enzymatic digestion was done at 37 °C overnight. The next day, digestion was quenched by addition of a final concentration of 2% TFA. Peptide concentration was then determined by a NanoDrop spectrophotometer (Thermo Fisher Scientific) and a total of 350 ng of peptides were loaded on Evotip C18 trap columns (Evosep Biosystems) according to the manufacturer’s instructions.

#### LC-MS/MS analysis

Peptides of hippocampus and gastrocnemius muscle samples were separated on 15-cm, 150-μM ID column packed with C18 beads (1.5 μm) (Pepsep) on an Evosep ONE HPLC system applying the default 30-SPD (30 samples per day) method. Column temperature was held at 50 ^0^C. Upon elution, peptides were injected via a CaptiveSpray source and 20-μm emitter into a timsTOF Pro 2 mass spectrometer (Bruker) operated in diaPASEF mode. MS data were collected over a 100-1700 m/z range. During MS/MS data collection each diaPASEF cycle was 1.8 seconds, covering the ion mobility range of 1.6-0.6 1/K0. Ion mobility was calibrated using three Agilent ESI-L Tuning Mix ions 622.0289, 922.0097 and 1221.9906. For diaPASEF we used a long-gradient method which included 10 diaPASEF scans with three 25 Da windows per ramp (for hippocampus) whereas 16 diaPASEF scans with two 25 Da windows per ramp (for muscle), mass range 400.0-1201.0 Da and mobility range 1.43-0.60 1/K_0_. The collision energy was decreased linearly from 59 eV at 1/K_0_ = 1.3 to 20 eV at 1/K_0_ = 0.85 Vs cm^-2^. Both accumulation time and PASEF ramp time was set to 100 ms. For the plasma proteomics, peptides were separated on a PepSep 8 cm, 150 μM ID reversed-phase column packed with C18 beads (1.5 μm) (PepSep) using an Evosep One LC system with the default 60-SDP (60 samples per day) method. Column temperature was maintained at 35 °C. Liquid chromatography was coupled via a CaptiveSpray ion source with a timsTOF Pro2 (Bruker Daltonics) operating in diaPASEF mode. MS data were collected over a 100-1700 m/z range. During each MS/MS data collection each dia-PASEF cycle was 0.95 seconds, covering the ion mobility range 1.4-0.6 1/K_0_. Ion mobility was calibrated using three Agilent ESI-L Tuning Mix ions 622.0289, 922.0097, and 1221.9906. For diaPASEF we used a modified version of the short-gradient method which included 8 diaPASEF scans with three 25-Da windows per ramp, mass range 400.0-1000.0 Da, and mobility range 1.37-0.64 1/K_0_. The collision energy was decreased linearly from 45 eV at 1/K_0_ = 1.3 to 27 eV at 1/K_0_ = 0.85 Vs cm-2. Both accumulation time and PASEF ramp time were set to 100 ms.

#### MS data analysis

Raw MS data was quantified using DIA-NN software v.1.8.1 [84]. The precursor mass and fragment mass were matched with an initial mass tolerance of 10 and 20 ppm, respectively. The search also included the following default parameters. The fixed modification of carbamidomethyl cysteine and N-terminal methionine excision were enabled. Peptide length was set to seven to 30 amino acids, and missed miscleavages were set to one. The precursor m/z range was set from 300 to 1800. The false discovery rate (FDR) was set to 1% at the peptide precursor level and protein level, and match between runs was enabled.

#### Differential proteome analysis

Proteomic data analysis was performed in R (v.4.3.0) with the following packages: clusterProfiler (v4.8.3), dplyr (v1.1.4), ggplot2 (v3.5.1), factoextra (v 1.0.7), limma (v3.56.2), PhosR (v1.10.0), plotly (v4.10.4), reshape2 (1.4.4), viridis (v0.6.5). Protein intensities were transformed to log2 scale, samples with low protein identifications (< 50 %) were removed, missing values were filtered to allow at least 60% of quantified values within each experimental group (WT-Con, WT-Ctsb, AD-Con and AD-Ctsb), and then quantile normalization was performed. In hippocampal and muscle proteomics, missing values were imputed from a normal distribution using tImpute function (width: 0.3, down shift: 1.8) by PhosR package. Subsequently, variance was corrected using weighted surrogate variance analysis (wsva) function from the limma package[85]. Differential expression analysis was done using lmFit function by limma package. Contrasts for different pairwise group comparisons, the main effects of CTSB treatment and AD pathogenesis as well as the interaction between CTSB and AD pathogenesis were assessed. Using Benjamini-Hochberg method, p values were corrected and FDR < 0.10 was considered as a cut-off for statistical significance. Gene set enrichment analyses were done by the ClusterProfiler package using biological process, cellular component and molecular function ontologies separately. Gene sets with adjusted p values < 0.05 were considered statistically significant.

### Statistical analysis

Statistical analyses were performed as described[86]. All behavioral and histological data was analyzed in RStudio (Version 1.4.1717 – 2009-2021 RStudio, PBC) R version 4.1.0 (2021-05-18) using either Generalized Linear Model (GLM) or Generalized Linear Mixed Model (GLMM). The family distribution and its canonical link function were chosen beforehand taking in account the nature of the data[87]. Specifically, continuous data with no boundaries were analyzed using Gaussian distribution with Identity link; continuous data presenting no upper boundaries but presenting lower boundaries = zero were analyzed using Gamma distribution with Log link; continuous data presenting both upper and lower boundaries were analyzed using Beta distribution with Logit link; discrete data (counts) presenting only lower boundaries = zero were first analyzed using Poisson distribution with Log link followed by Negative Binomial distribution with Log link for the cases in which over-dispersion was detected during diagnosis procedure. GLM and GLMM were performed using either lme4 package[88] or glmmTMB package [89]. When necessary, the raw data were subjected to transformations before the analysis, as described for each analysis in the Supplementary Material.

A backward stepwise selection model approach (based on the Akaike Information Criterion – AIC – through the use of the stepAIC function) or, alternatively, an approach evaluating all possible combinations of subsets of factors (based on the Akaike Information Criterion corrected for small samples – AICc – through the use of the dredge function from the MuMin package) was applied to select the most parsimonious model. The model presenting the lowest AIC (or AICc) value was selected. All selected models were submitted to diagnosis using the DHARMa package[90]. Except for overdispersion in the models using Poisson distribution (as described above), no action was made to avoid diagnosis problems (diagnosis outputs are described for each individual variable in Supplementary Material). Estimated marginal means and 95% CI were extracted from the selected models using emmeans package[91] and were graph plotted using GraphPad Prism 8. Bonferroni pairwise (post hoc) comparisons were performed for the cases in which any interaction effect was detected, controlling for family-wise error. For time spent in target quadrant data (probe trials in the MWM paradigm), an additional analysis was performed in which each group (WT-Con, WT-Ctsb, AD-Con, and AD-Ctsb) was individually compared against the 25% chance of randomly staying in each of the quadrants (One-Sample Wilcoxon Signed Rank Test). Correlations were evaluated by Spearman’s rank correlation. A 0.05 significance level (alpha) was set for all analyses. All the statistical results are presented in detail in the Supplementary Tables.

### Bias-reducing measures

Randomization, blinding and systematic sampling procedures were applied to minimize both performance and detection bias for histological data. Detection bias was further minimized by using automated systems (equivalent to blinding outcome assessors) to collect behavioral data. Attrition bias was minimized by adhering to predefined exclusion criteria as follows: (i) identification of signs of pain or distress in the mice (no animals were excluded based on this criterion, however, one AD-Ctsb mouse died between the last behavioral testing session and euthanasia). (ii) Poor staining/image quality resulted in exclusion of 1 animal (WT-Ctsb) in DCX analysis; 2 animals (1 WT-Con and 1 AD-Ctsb) in GFAP/Iba1 analysis; 3 AD-Ctsb mice in ThioS analysis. No outliers were excluded. Based on “ARRIVE guidelines 2.0: Essential 10 list”[92] available at https://arriveguidelines.org/arrive-guidelines and on “SYRCLE’s risk of bias tool for animal studies”[93] all efforts were made to ensure transparency and avoid reporting bias[86].

## DATA AVAILABILITY

The mass spectrometry proteomics data have been deposited in the ProteomeXchange Consortium (http://proteomecentral.proteomexchange.org) via the PRIDE partner repository with the dataset identifier PXD057069.

## ACKNOWLEDGEMENTS

We thank the FAU NeuroBehavioral, NeuroImaging and Comparative Medicine Cores for assistance, Alcira Munchow and Drs. Jana Boerner, Daniel Nemeth and Carmen Vivar for technical advice, Stephanie Vitek for graphic art contributions and Linda R. Kitabayashi for photomicrograph composition. Mass spectrometry analyses were performed by the Proteomics Research Infrastructure (PRI) at the University of Copenhagen (UCPH), supported by the Novo Nordisk Foundation (NNF) (grant agreement number NNF19SA0059305). The work was also supported by grants from Novo Nordisk Foundation (grant agreement number NNF23SA0084103, NNF18CC0034900), NIH/NHLBI grant RO1HL155986 to T.K., and by the Ed and Ethel Moore Alzheimer’s Disease Research Program of the Florida Department of Health (9AZ02 to H.v.P.).

## AUTHOR CONTRIBUTIONS

A.P., H.H., C.M.L., A.E., C.A., O.C., A.W., G.M., K.S., T.K., performed experiments. A.P., C.M.L., H.H., A.A. performed data analysis. Methodology: S.G., T.K. Conceptualization: H.v.P. Writing, original draft: A.P., H.H., C.M.L., A.S.D., H.v.P. Writing, review and editing: A.A., R.B. and T.K. Supervision: A.S.D., R.B., H.v.P.

