## Supplementary Figures and Tables for "Muscle Cathepsin B treatment improves behavioral and neurogenic deficits in a mouse model of Alzheimer’s Disease"

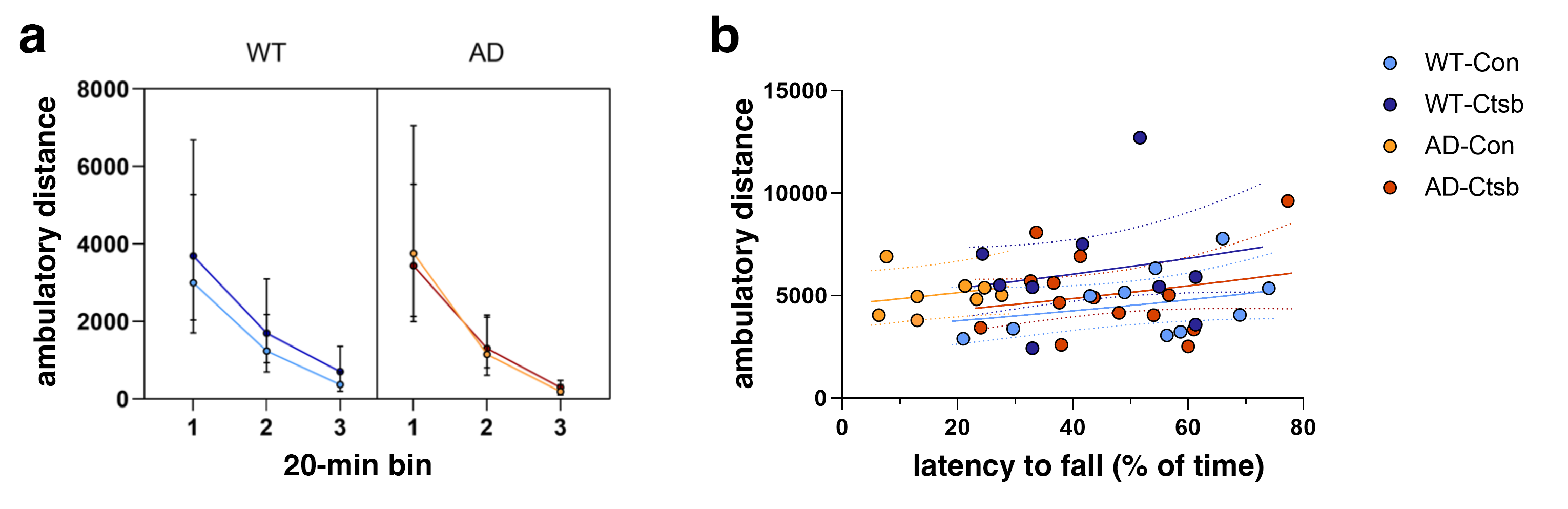

**A**

**B**

**Figure S1.** **Activity Box.** **(A)** Distance traveled in the activity box (open field) is increased in WT-Ctsb mice (Table S1). Analysis of the ambulatory distance over time (20 min bins) revealed a Time effect (GLMM analysis) in which Bin 1 > Bin 2 > Bin 3 for all groups (Table S2). This suggests that all groups habituated similarly to the arena and that between groups differences in total ambulatory distance are a result of distinct activity levels. **(B)** Generalized Linear Model (GLM) analysis revealed an interaction between genotype and treatment, and a latency to fall effect that indicates a within-group relationship in which mice that fall sooner in the rotarod traveled a shorter distance than mice that fall later (Table S1). Solid lines represent the Estimated Marginal Means while dashed lines represent 95% CI. (N; WT-Con = 10, WT-Ctsb = 9, AD-Con = 8, AD-Ctsb = 14). Data are presented as Estimated Marginal Means and their respective 95% CI.

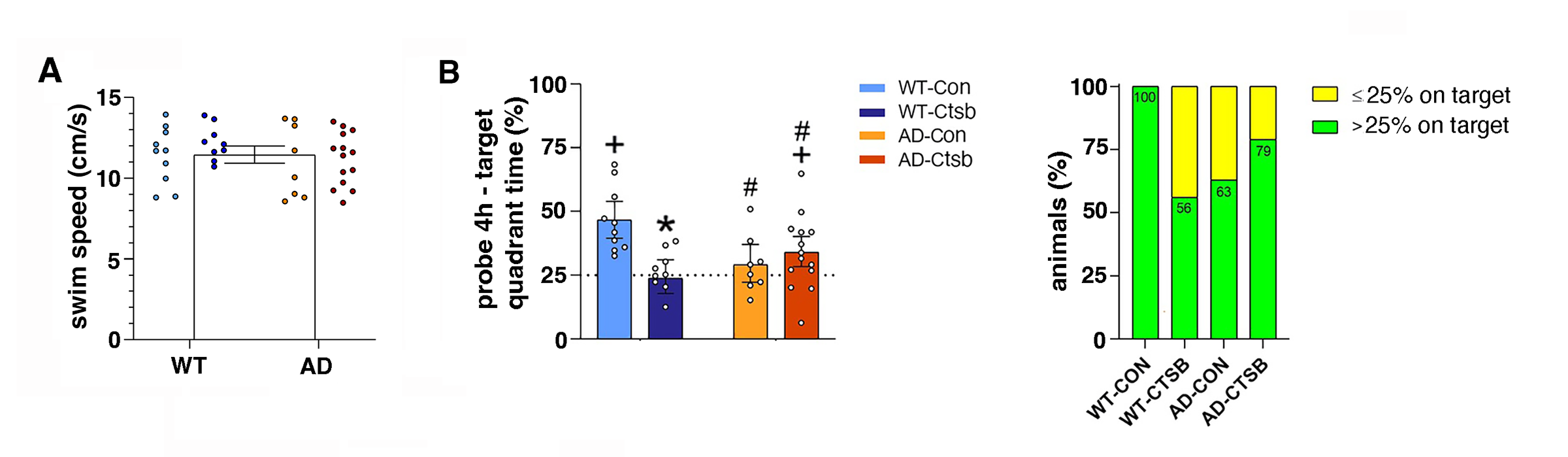

Figure S2. Morris water maze performance (A) Swim speed. Average swim speed did not differ between the groups on acquisition day 1 (Table S5) and did not change over time (Table S6). The single bar and whiskers represent the Estimated Marginal Means and its 95% CI (respectively) estimated by the model for the entire data set (i.e., all groups pooled together). (B) Target quadrant time in the 4 h probe trial. In the 4h probe trial the WT-Con group spent more time in the target quadrant than the WT-Ctsb and AD-Con groups. AD-Ctsb mice spent more time in the target quadrant than WT-Ctsb mice. Indeed, both the WT-Con and AD-Ctsb groups preferred the target quadrant above 25% chance, indicating that these groups remember the location of the platform that was present during the acquisition phase. Specifically, WT-Con and AD-Ctsb mice were in the target quadrant 46% (p<0.001) and 34% (p=0.024), respectively, of the time, whereas the AD-Con mice and WT-Ctsb spent 29% (p=0.19) and 23% (p=0.36) of the time, respectively, and as depicted in the contingency graph of the percentage of mice that spent either more (in green) or less (in yellow) than 25% of the time in the target quadrant (Table S8). (N; WT-Con = 10, WT-Ctsb = 9, AD-Con = 8, AD-Ctsb = 14). Data were analyzed by GLM and are presented as Estimated Marginal Means and their respective 95% CI. # P<0.05 AD compared to WT in the same Treatment. *P<0.05 Ctsb compared to Con in the same Genotype. ^+^P<0.05 compared to 25% chance (one-tailed Wilcoxon signed rank exact test).

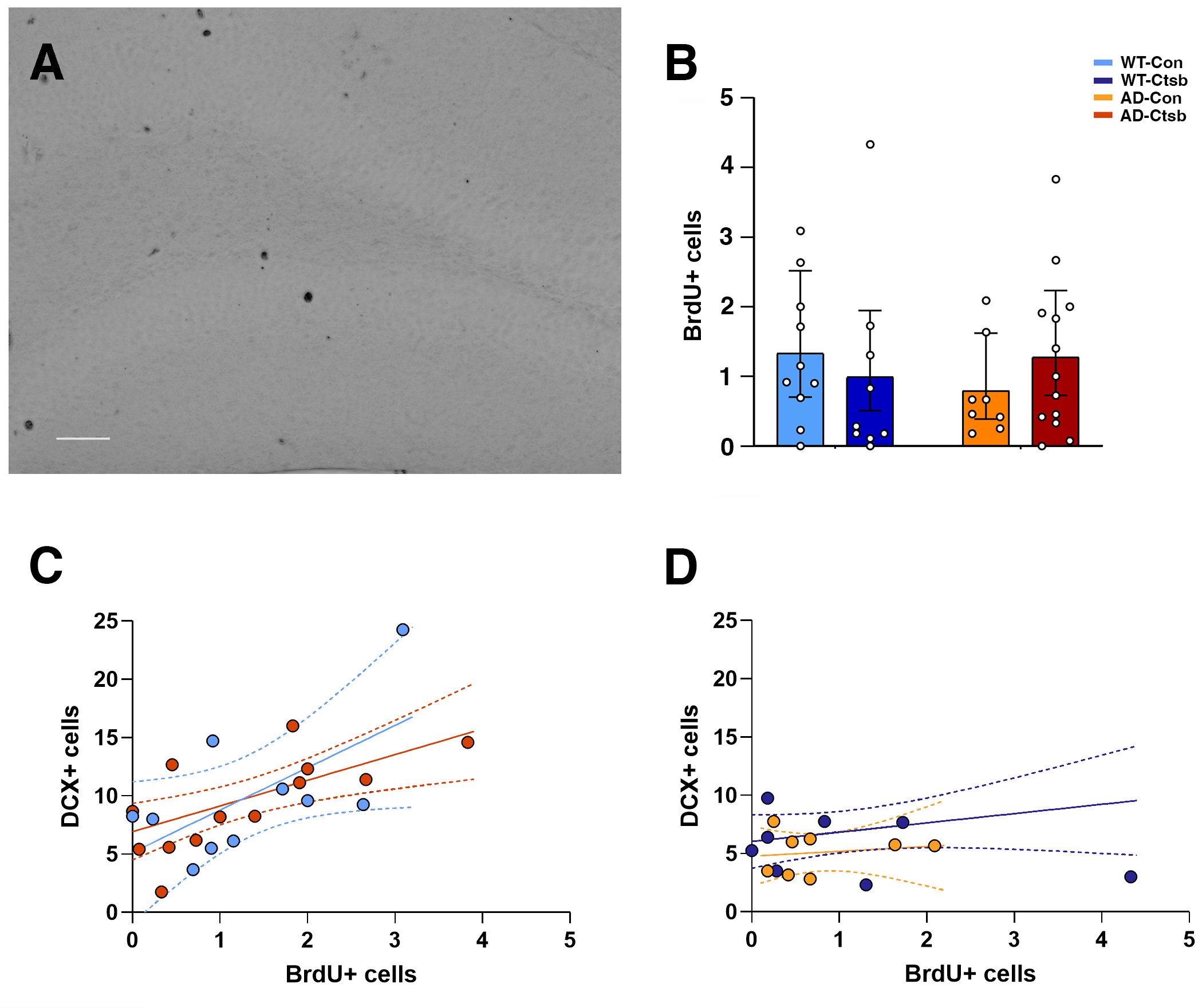

Figure S3. Long-term adult-born cell survival (A) Representative photomicrograph of BrdU labeling in the dentate gyrus. Scale bar 50 μm. (B) BrdU^+^ cells per section did not differ between the groups (mean = 1.13; 95% CI 0.84 – 1.54; Table S15). (C,D) Spearman correlation analysis between DCX^+^ cells and BrdU^+^ cells (Table S16). (C) BrdU^+^ and DCX^+^ cell number increased concurrently in both WT-Con (P=0.0665) and AD-Ctsb (P= 0.0183) groups, but not in (D) WT-Ctsb (P= 0.35) and AD-Con (P= 0.91) groups. In B, data were analyzed by GLM and are presented as Estimated Marginal Means and their respective 95% CI. In C and D, solid lines represent the correlation for each group while dashed lines represent their respective 95% CI.

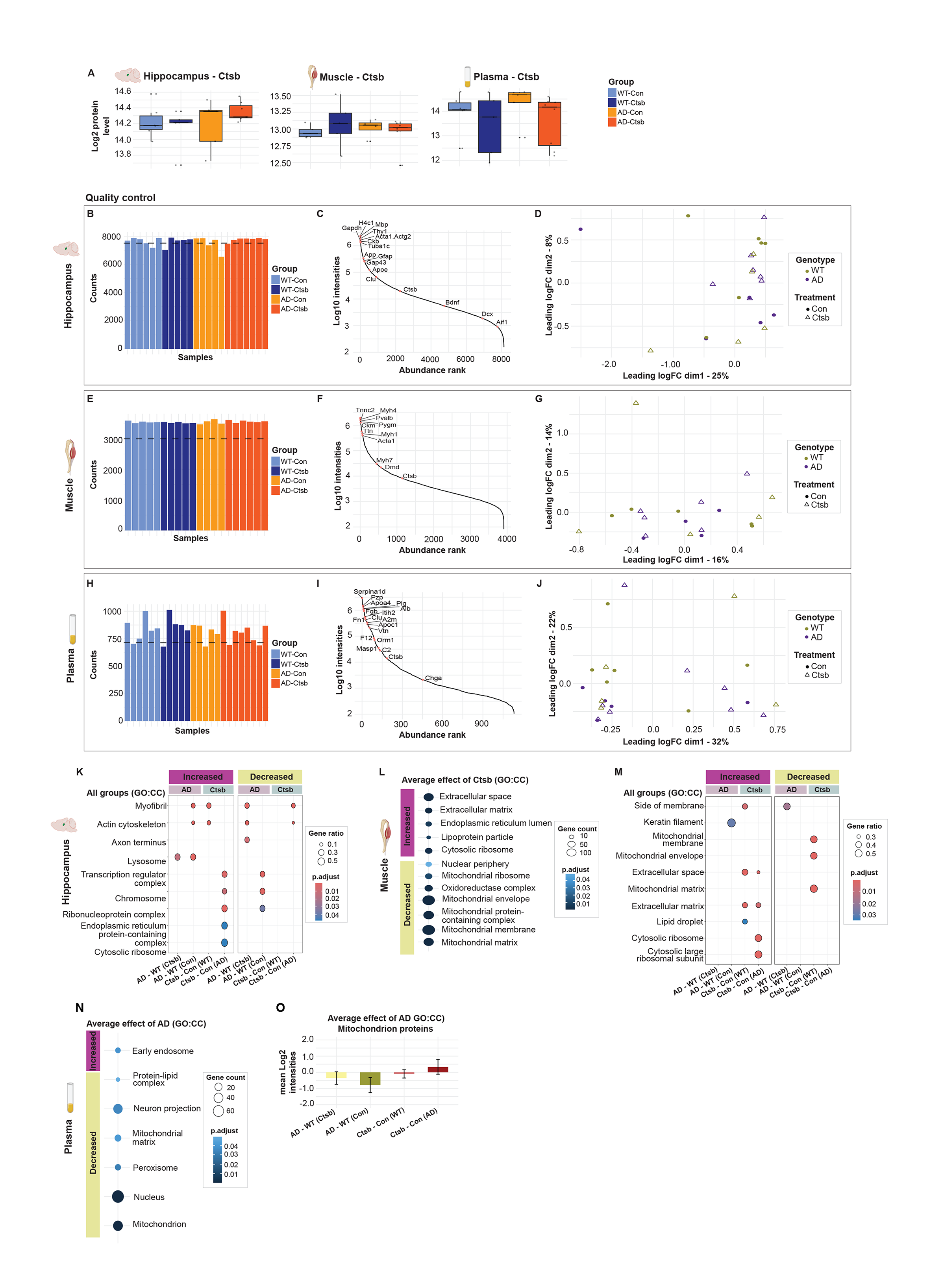
Figure S4. Additional proteomics data, quality control and gene set enrichment analyses.

(A) Protein abundance of Ctsb in hippocampus, muscle and plasma across all experimental groups. Quality control of (B-D) hippocampal, (E-G) muscle and (H-J) plasma proteomics. After filtering, (B) 7509 proteins were reliably quantified in the hippocampus, (E) 2998 proteins in the muscle and (H) 713 proteins in the plasma. Dynamic range was over four orders of magnitude and proteins of interest were marked in the (C) hippocampus, (F) muscle and (I) plasma. Multidimensional scaling did not show any indication of genotype or treatment-specific clustering in the (D) hippocampus, (G) muscle and (J) plasma. (K) GSEA of the hippocampus, summarizing enriched cellular component (CC) ontologies in all groups. (L-M) GSEA of the muscle, showing enriched CC ontologies (l) by average effect of Ctsb treatment and (M) across all groups. (N) GSEA of the plasma, showing enriched CC terms based on the average effect of AD. (O) Mean abundance of proteins related to mitochondrion showed a tendency towards increase only in AD-Ctsb plasma. For GSEAs, a cut-off of FDR < 0.05 was used. (K,M,O) Data for pairwise comparisons are annotated as follows: AD-WT(Ctsb), *AD-Ctsb vs WT-Ctsb*; AD-WT(Con), *AD-Con vs WT-Con*; Ctsb-Con(WT), *WT-Ctsb vs WT-Con*; Ctsb-Con(AD), *AD-Ctsb vs AD-Con*.

Table S1. Summary of the selected model for analysis of total distance traveled during 60 min session in activity box

| **Method** | Generalized linear model (GLM) | | | | |
| --- | --- | --- | --- | --- | --- |
| **Family distribution with link function** | Gamma ( log ) | | | | |
| **Formula** | Total distance traveled ~ Genotype * Treatment + Latency to fall | | | | |
| **Fixed effects** |  | **β** | **SE** | **t value** | **Pr(>\|t\|)** |
|  | Intercept | 8.1152 | 0.2479 | 32.740 | <2e-16 *** |
|  | GenAD | 0.3103 | 0.2266 | 1.370 | 0.1793 |
|  | TreatCTSB | 0.3522 | 0.1707 | 2.063 | 0.0464 * |
|  | Latency to fall | 0.5995 | 0.4218 | 1.421 | 0.1639 |
|  | GenAD:TreatCTSB | -0.5297 | 0.2811 | -1.884 | 0.0676 |
| **Diagnosis** | No diagnosis problems detected | | | | |
| **Other information** | Dispersion parameter for Gamma family taken to be 0.1313971  Null deviance: 5.4227 on 40 degrees of freedom  Residual deviance: 4.7259 on 36 degrees of freedom | | | | |

β = estimated coefficients for the fixed effect terms; SE = Standard Error; Intercept = WT-CON; Significance codes: *** 0.001; ** 0.01; * 0.05

Table S2. Summary of the selected model for analysis of distance traveled over time (20 min bins) during 60 min session in activity box

| **Method** | Generalized linear mixed model (GLMM) | | | | | |
| --- | --- | --- | --- | --- | --- | --- |
| **Family distribution with link function** | Gamma ( log ) | | | | | |
| **Formula** | Distance traveled ~ Time + (1 \| id) | | | | | |
| **Random effects** |  | **Variance** | **SD** | |  |  |
|  | id | 0.09638 | 0.3104 | |  |  |
| **# of observations** | 123 |  |  | |  |  |
| **# of id** | 41 |  |  | |  |  |
| **Fixed effects** |  | **β** | **SE** | **z value** | | **Pr(>\|z\|)** |
|  | Intercept | 8.1621 | 0.1481 | 55.09 | | < 2e-16 *** |
|  | Bin2 | -0.9795 | 0.1987 | -4.93 | | 8.23e-07 *** |
|  | Bin3 | -2.3121 | 0.2172 | -10.64 | | <2e-16 *** |
| **Diagnosis** | Residuals normality problems detected;  Residuals dispersion problems detected;  Outliers detected;  Within-group deviations from uniformity detected; | | | | | |
| **Other information** | Before starting the analysis, the data were corrected for the use of Gamma distribution through the following procedure:  A positive constant (epsilon = 0.000001) was add to each individual value  Models ranked by AICc (dredge function from the MuMin package)  Dispersion parameter for Gamma family taken to be 0.782  Estimated Marginal Means and their respective 95% Confidence Intervals  (values are presented as cm)  Time:  Bin 1 = 3505; 2614 – 4701  Bin 2 = 1316; 981 – 1766 ^a^  Bin 3 = 347; 250 – 482 ^a,b^ | | | | | |

SD = Standard Deviation; β = estimated coefficients for the fixed effect terms; SE = Standard Error; Intercept = Bin 1; ^a^ indicates p= 0.0001 when compared to Bin 1; ^b^ indicates p= 0.0001 when compared to Bin 2; Significance codes: *** 0.001; ** 0.01; * 0.05

**Table S3: Summary of the selected model for analysis of latency to fall in the rotarod test**

| **Method** | Generalized linear model (GLM) | | | | |
| --- | --- | --- | --- | --- | --- |
| **Family distribution with link function** | Beta ( logit ) | | | | |
| **Formula** | Latency to fall ~ Genotype * Treatment | | | | |
| **Fixed effects** |  | **β** | **SE** | **z value** | **Pr(>\|z\|)** |
|  | Intercept | 0.07165 | 0.17334 | 0.413 | 0.679 |
|  | GenAD | -1.57622 | 0.29597 | -5.326 | 1.01e-07 *** |
|  | TreatCTSB | -0.34508 | 0.25288 | -1.365 | 0.172 |
|  | GenAD:TreatCTSB | 1.70036 | 0.37839 | 4.494 | 7.00e-06 *** |
| **Diagnosis** | No diagnosis problems detected | | | | |
| **Other information** | Before starting the analysis, the data were corrected for the use of Beta distribution through the following procedure:  Each latency to fall value (y) was obtaining as a frequency of the total duration of the test (i.e., 0 < y < 1), as follows:  Latency to fall (s) / 300 (s);  Dispersion parameter for Beta family taken to be 12.3 | | | | |

β = estimated coefficients for the fixed effect terms; SE = Standard Error; Intercept = WT-CON; Significance codes: *** 0.001; ** 0.01; * 0.05

**Table S4: Summary of the selected model for analysis of latency to find the platform during the acquisition phase (4-day bins) of the Morris water maze paradigm**

| **Method** | | Generalized linear mixed model (GLMM) | | | | | |
| --- | --- | --- | --- | --- | --- | --- | --- |
| **Family distribution with link function** | | Beta ( logit ) | | | | | |
| **Formula** | | Latency to find the platform ~ Genotype * Treatment * Time + (1 \| id) | | | | | |
| **Random effects** | |  | **Variance** | **SD** | |  |  |
|  | | id | 0.5702 | 0.7551 | |  |  |
| **# of observations** | | 205 |  |  | |  |  |
| **# of id** | | 41 |  |  | |  |  |
| **Fixed effects** |  | | **β** | **SE** | **z value** | | **Pr(>\|z\|)** |
|  | Intercept | | 2.65612 | 0.38324 | 6.931 | | 4.19e-12 *** |
|  | GenAD | | 1.48835 | 0.60539 | 2.458 | | 0.013953 * |
|  | TreatCTSB | | -0.14407 | 0.54371 | -0.265 | | 0.791021 |
|  | Bin2 | | -2.42801 | 0.35810 | -6.780 | | 1.20e-11 *** |
|  | Bin3 | | -3.43976 | 0.36732 | -9.365 | | < 2e-16 *** |
|  | Bin4 | | -3.29858 | 0.36579 | -9.018 | | < 2e-16 *** |
|  | Bin5 | | -3.97430 | 0.37888 | -10.490 | | < 2e-16 *** |
|  | GenAD:TreatCTSB | | -1.17818 | 0.78560 | -1.500 | | 0.133685 |
|  | GenAD:Bin2 | | -1.03438 | 0.57398 | -1.802 | | 0.071527 |
|  | GenAD:Bin3 | | 0.29789 | 0.58088 | 0.513 | | 0.608078 |
|  | GenAD:Bin4 | | -0.65492 | 0.57451 | -1.140 | | 0.254302 |
|  | GenAD:Bin5 | | -0.61197 | 0.59110 | -1.035 | | 0.300525 |
|  | TreatCTSB:TimeBin2 | | -0.16939 | 0.50542 | -0.335 | | 0.737518 |
|  | TreatCTSB:TimeBin3 | | 0.10379 | 0.51927 | 0.200 | | 0.841583 |
|  | TreatCTSB:TimeBin4 | | 0.27863 | 0.51889 | 0.537 | | 0.591286 |
|  | TreatCTSB:TimeBin5 | | -0.01328 | 0.53588 | -0.025 | | 0.980225 |
|  | GenAD:TreatCTSB:TimeBin2 | | 2.96463 | 0.76328 | 3.884 | | 0.00010 *** |
|  | GenAD:TreatCTSB:TimeBin3 | | 0.12509 | 0.74823 | 0.167 | | 0.867230 |
|  | GenAD:TreatCTSB:TimeBin4 | | 1.05775 | 0.75049 | 1.409 | | 0.158713 |
|  | GenAD:TreatCTSB:TimeBin5 | | 1.45907 | 0.76984 | 1.895 | | 0.058053 |
| **Diagnosis** | Residuals normality problems detected;  Residuals dispersion problems detected;  Quantile deviations detected; | | | | | | |
| **Other information** | Before starting the analysis, the data were corrected for the use of Beta distribution through the following procedure:  For each 4-day bin, each latency to find the platform value (y) was obtaining as a frequency of the total duration of the test (i.e., 0 < y < 1), as follows:  Latency to find the platform (s) / 60 (s);  Later, a negative constant (epsilon = -0.000001) was add to each individual value  Dispersion parameter for Beta family taken to be 9.67 | | | | | | |

SD = Standard Deviation; β = estimated coefficients for the fixed effect terms; SE = Standard Error; Intercept = WT-CON Bin 1; Significance codes: *** 0.001; ** 0.01; * 0.05

**Table S5: Summary of the selected model for analysis of average speed during day 1 of the acquisition phase of the Morris water maze paradigm**

| **Method** | Generalized linear model (GLM) | | | | |
| --- | --- | --- | --- | --- | --- |
| **Family distribution with link function** | Gamma ( log ) | | | | |
| **Formula** | Average speed day 1 ~ 1 | | | | |
| **Fixed effects** |  | **β** | **SE** | **t value** | **Pr(>\|t\|)** |
|  | Intercept | 2.43768 | 0.02302 | 105.9 | <2e-16 *** |
| **Diagnosis** | No diagnosis problems detected | | | | |
| **Other information** | Dispersion parameter for Gamma family taken to be 0.02172645  Null deviance: 0.91201 on 40 degrees of freedom  Residual deviance: 0.91201 on 40 degrees of freedom  Estimated Marginal Means and their respective 95% Confidence Intervals for the full model, i.e., Average speed day 1 ~ Genotype * Treatment  (values are presented as cm/s)  Treatment = CON:  WT = 11.4; 10.4 – 12.6  AD = 11.1; 9.96 – 12.4  Treatment = CTSB:  WT = 12.2; 11.0 – 13.5  AD = 11.2; 10.32 – 12.1 | | | | |

β = estimated coefficients for the fixed effect terms; SE = Standard Error; Intercept = Overall; Significance codes: *** 0.001; ** 0.01; * 0.05

**Table S6: Summary of the selected model for analysis of average speed during the acquisition phase (4-day bins) of the Morris water maze paradigm**

| **Method** | | Generalized linear mixed model (GLMM) - Linear mixed model fit by maximum likelihood ['lmerMod'] | | | |
| --- | --- | --- | --- | --- | --- |
| **Family distribution with link function** | | Gaussian ( identity ) | | | |
| **Formula** | | Average speed ~ 1 + (1 \| id) | | | |
| **Random effects** | |  | **Variance** | **SD** | |
|  | | id | 0.03044 | 0.1745 | |
|  | | Residual | 0.56794 | 0.7536 | |
| **# of observations** | | 205 |  |  | |
| **# of id** | | 41 |  |  | |
| **Fixed effects** |  | | **β** | **SE** | **t value** |
|  | Intercept | | 11.33932 | 0.05927 | 191.3 |
| **Diagnosis** | No diagnosis problems detected | | | | |
| **Other information** | GLMM did not run when using Gamma distribution ( log ). Because the variable is continuous and present no values near to the lower boundary, the model was run using Gaussian distribution ( identity ).  Models ranked by AICc (dredge function from the MuMin package) | | | | |

SD = Standard Deviation; β = estimated coefficients for the fixed effect terms; SE = Standard Error; Intercept = Overall; Significance codes: *** 0.001; ** 0.01; * 0.05

**Table S7: Summary of the selected model for analysis of time spent in the target quadrant during the 4h probe trial in the Morris water maze**

| **Method** | Generalized linear model (GLM) | | | | |
| --- | --- | --- | --- | --- | --- |
| **Family distribution with link function** | Beta ( logit ) | | | | |
| **Formula** | Time spent in the target quadrant ~ Genotype + Treatment + Speed + Genotype:Treatment + Treatment:Speed | | | | |
| **Fixed effects** |  | **β** | **SE** | **z value** | **Pr(>\|z\|)** |
|  | Intercept | -0.069466 | 0.690288 | -0.101 | 0.919841 |
|  | GenAD | -0.755924 | 0.227027 | -3.330 | 0.000869 *** |
|  | TreatCTSB | -3.772909 | 1.167028 | -3.233 | 0.001225 ** |
|  | Speed | -0.005502 | 0.059120 | -0.093 | 0.925856 |
|  | GenAD:TreatCTSB | 1.256673 | 0.317114 | 3.963 | 7.41e-05 *** |
|  | TreatCTSB:Speed | 0.239705 | 0.095948 | 2.498 | 0.012480 * |
| **Diagnosis** | No diagnosis problems detected | | | | |
| **Other information** | Before starting the analysis, the data were corrected for the use of Beta distribution through the following procedure:  Each time spent in the target quadrant (y) was obtaining as a frequency of the total duration of the test (i.e., 0 < y < 1), as follows:  Time spent in the target quadrant (s) / 60 (s);  Dispersion parameter for Beta family taken to be 18.5  Estimated Marginal Means and their respective 95% Confidence Intervals when Speed is fixed at 11.4 cm/s (the overall average speed during day 1)  (values are presented as %)  Treatment = CON:  WT = 46.7; 39.6 – 54.0  AD = 29.1; 22.3 – 37.1  Treatment = CTSB:  WT = 23.8; 17.9 – 31.0  AD = 34.1; 28.4 – 40.2 | | | | |

Speed = Average speed during day 1 of the acquisition phase; β = estimated coefficients for the fixed effect terms; SE = Standard Error; Intercept = WT-CON; Significance codes: *** 0.001; ** 0.01; * 0.05

**Table S8: Summary of the one-sample Wilcoxon signed rank test for analysis of time spent in the target quadrant during the 4h probe trial in the Morris water maze paradigm against the 25% chance**

| **Method** | One-sample Wilcoxon signed rank exact test (one-tail test) | | | | |
| --- | --- | --- | --- | --- | --- |
| **Reference value** | 25% (mu = 0.25) | | | | |
| **Alternative Hypothesis** | one-tail test where the H1 = location is greater than 25% chance | | | | |
| **Output** |  | **V value** | **N** | **p-value** | **one-tail** |
|  | WT-CON | 55 | 10 | 0.000977 | greater |
|  | WT-CTSB | 26 | 9 | 0.3672 | equal |
|  | AD-CON | 25 | 8 | 0.1914 | equal |
|  | AD-CTSB | 84 | 14 | 0.02472 | greater |

N= sample size

**Table S9: Summary of the selected model for analysis of time spent in the target quadrant during the 24h probe trial in the Morris water maze**

| **Method** | Generalized linear model (GLM) | | | | |
| --- | --- | --- | --- | --- | --- |
| **Family distribution with link function** | Beta ( logit ) | | | | |
| **Formula** | Time spent in the target quadrant ~ Genotype * Treatment * Speed | | | | |
| **Fixed effects** |  | **β** | **SE** | **z value** | **Pr(>\|z\|)** |
|  | Intercept | -0.72510 | 0.91886 | -0.789 | 0.430038 |
|  | GenAD | -4.00382 | 1.41351 | -2.833 | 0.004618 ** |
|  | TreatCTSB | -4.54110 | 2.10162 | -2.161 | 0.030714 * |
|  | Speed | 0.02693 | 0.07961 | 0.338 | 0.735174 |
|  | GenAD:TreatCTSB | 8.64317 | 2.51605 | 3.435 | 0.000592 *** |
|  | GenAD:Speed | 0.25822 | 0.12030 | 2.146 | 0.031842 * |
|  | TreatCTSB:Speed | 0.32570 | 0.17222 | 1.891 | 0.058601 |
|  | GenAD:TreatCTSB:Speed | -0.62992 | 0.20913 | -3.012 | 0.002595 ** |
| **Diagnosis** | No diagnosis problems detected | | | | |
| **Other information** | Before starting the analysis, the data were corrected for the use of Beta distribution through the following procedure:  Each time spent in the target quadrant (y) was obtaining as a frequency of the total duration of the test (i.e., 0 < y < 1), as follows:  Time spent in the target quadrant (s) / 60 (s);  Dispersion parameter for Beta family taken to be 22.2  Estimated Marginal Means and their respective 95% Confidence Intervals when Speed is fixed at 11.4 cm/s (the overall average speed during day 1)  (values are presented as %)  Treatment = CON:  WT = 39.7; 33.4 – 46.4  AD = 18.8; 13.6 – 25.3  Treatment = CTSB:  WT = 22.6; 16.1 – 30.8  AD = 30.0; 25.1 – 35.5 | | | | |

Speed = Average speed during day 1 of the acquisition phase; β = estimated coefficients for the fixed effect terms; SE = Standard Error; Intercept = WT-CON; Significance codes: *** 0.001; ** 0.01; * 0.05

**Table S10: Summary of the one-sample Wilcoxon signed rank test for analysis of time spent in the target quadrant during the 24h probe trial in the Morris water maze paradigm against the 25% chance**

| **Method** | One-sample Wilcoxon signed rank exact test (one-tail test) | | | | |
| --- | --- | --- | --- | --- | --- |
| **Reference value** | 25% (mu = 0.25) | | | | |
| **Alternative Hypothesis** | one-tail test where the H1 = location is greater than 25% chance | | | | |
| **Output** |  | **V value** | **N** | **p-value** | **one-tail** |
|  | WT-CON | 55 | 10 | 0.000977 | greater |
|  | WT-CTSB | 29 | 9 | 0.248 | equal |
|  | AD-CON | 14 | 8 | 0.7266 | equal |
|  | AD-CTSB | 81 | 14 | 0.03925 | greater |

N= sample size

**Table S11: Summary of the selected model for analysis of percentage of freezing during the conditioning phase of the fear conditioning paradigm**

| **Method** | | Generalized linear mixed model (GLMM) | | | | | |
| --- | --- | --- | --- | --- | --- | --- | --- |
| **Family distribution with link function** | | Beta ( logit ) | | | | | |
| **Formula** | | Freezing behavior ~ Genotype * Treatment * Tone-Shock + (1 \| id) | | | | | |
| **Random effects** | |  | **Variance** | **SD** | |  |  |
|  | | id | 0.2987 | 0.5465 | |  |  |
| **# of observations** | | 123 |  |  | |  |  |
| **# of id** | | 41 |  |  | |  |  |
| **Fixed effects** |  | | **β** | **SE** | **z value** | | **Pr(>\|z\|)** |
|  | Intercept | | -0.6223 | 0.3731 | -1.668 | | 0.09532 |
|  | GenAD | | 0.4815 | 0.5490 | 0.877 | | 0.38044 |
|  | TreatCTSB | | -0.4615 | 0.5412 | -0.853 | | 0.39380 |
|  | TS2 | | 1.1367 | 0.4559 | 2.493 | | 0.01265 * |
|  | TS3 | | 1.5231 | 0.4631 | 3.288 | | 0.00101 ** |
|  | GenAD:TreatCTSB | | 0.1192 | 0.7427 | 0.160 | | 0.87250 |
|  | GenAD:TS2 | | -0.7094 | 0.6700 | -1.059 | | 0.28969 |
|  | GenAD:TS3 | | -0.6574 | 0.6675 | -0.985 | | 0.32470 |
|  | TreatCTSB:TimeTS2 | | -0.8111 | 0.6592 | -1.230 | | 0.21854 |
|  | TreatCTSB:TimeTS3 | | -0.8471 | 0.6519 | -1.299 | | 0.19382 |
|  | GenAD:TreatCTSB:TimeTS2 | | 0.8975 | 0.9062 | 0.990 | | 0.32198 |
|  | GenAD:TreatCTSB:TimeTS3 | | 1.1078 | 0.8938 | 1.239 | | 0.21522 |
| **Diagnosis** | No diagnosis problems detected | | | | | | |
| **Other information** | Before starting the analysis, the data were corrected for the use of Beta distribution through the following procedure:  For each Tone-Shock presentation, each freezing behavior value (y) was obtaining as a frequency of the total duration of each tone presentation (i.e., 0 < y < 1), as follows:  Freezing behavior (s) / 30 (s);  Dispersion parameter for Beta family taken to be 3.18 | | | | | | |

SD = Standard Deviation; β = estimated coefficients for the fixed effect terms; SE = Standard Error; Intercept = WT-CON Tone-Shock presentation 1; TS = Tone-Shock presentation; Significance codes: *** 0.001; ** 0.01; * 0.05

**Table S12: Summary of the selected model for analysis of percentage of freezing during the tone-cued phase of the fear conditioning paradigm**

| **Method** | Generalized linear model (GLM) | | | | |
| --- | --- | --- | --- | --- | --- |
| **Family distribution with link function** | Beta ( logit ) | | | | |
| **Formula** | Freezing behavior ~ Genotype + Treatment + TS1 + Genotype:Treatment +  Genotype:TS1 + Treatment:TS1 | | | | |
| **Fixed effects** |  | **β** | **SE** | **z value** | **Pr(>\|z\|)** |
|  | Intercept | 0.606080 | 0.750491 | 0.808 | 0.4193 |
|  | GenAD | -1.287511 | 0.954155 | -1.349 | 0.1772 |
|  | TreatCTSB | 0.003667 | 0.795825 | 0.005 | 0.9963 |
|  | TS1 | 0.009905 | 0.021068 | 0.470 | 0.6382 |
|  | GenAD:TreatCTSB | 1.267221 | 0.813075 | 1.559 | 0.1191 |
|  | GenAD:TS1 | 0.033867 | 0.020073 | 1.687 | 0.0916 |
|  | TreatCTSB:TS1 | -0.034950 | 0.020324 | -1.720 | 0.0855 |
| **Diagnosis** | No diagnosis problems detected | | | | |
| **Other information** | Before starting the analysis, the data were corrected for the use of Beta distribution through the following procedure:  For each Tone presentation (T1-T3), each freezing behavior value (y) was obtaining as a frequency of the total duration of each tone presentation (i.e., 0 < y < 1), as follows: Freezing behavior (s) / 30 (s);  After that, the average frequency for T1-T3 was calculated. Later, a negative constant (epsilon = -0.000001) was add to each individual value  Dispersion parameter for Beta family taken to be 1.8 | | | | |

SD = Standard Deviation; β = estimated coefficients for the fixed effect terms; SE = Standard Error; Intercept = WT-CON Tone presentation 1; TS1 = First Tone-Shock presentation during the conditioning phase; T = Tone presentation; Significance codes: *** 0.001; ** 0.01; * 0.05

**Table S13: Summary of the selected model for analysis of percentage of freezing during the contextual phase of the fear conditioning paradigm**

| **Method** | | Generalized linear mixed model (GLMM) | | | | | |
| --- | --- | --- | --- | --- | --- | --- | --- |
| **Family distribution with link function** | | Beta ( logit ) | | | | | |
| **Formula** | | Freezing behavior ~ Genotype * Treatment * Session + (1 \| id) | | | | | |
| **Random effects** | |  | **Variance** | **SD** | |  |  |
|  | | id | 0.5312 | 0.7288 | |  |  |
| **# of observations** | | 164 |  |  | |  |  |
| **# of id** | | 41 |  |  | |  |  |
| **Fixed effects** |  | | **β** | **SE** | **z value** | | **Pr(>\|z\|)** |
|  | Intercept | | -1.93143 | 0.38417 | -5.028 | | 4.97e-07 *** |
|  | GenAD | | -0.30933 | 0.59103 | -0.523 | | 0.60071 |
|  | TreatCTSB | | -0.28857 | 0.56778 | -0.508 | | 0.61129 |
|  | Session2 | | 0.55537 | 0.40801 | 1.361 | | 0.17346 |
|  | Session3 | | 0.69847 | 0.40197 | 1.738 | | 0.08228 |
|  | Session4 | | 0.48018 | 0.40974 | 1.172 | | 0.24124 |
|  | GenAD:TreatCTSB | | 0.54790 | 0.79966 | 0.685 | | 0.49324 |
|  | GenAD:Session2 | | 2.70447 | 0.62860 | 4.302 | | 1.69e-05 *** |
|  | GenAD:Session3 | | 1.54914 | 0.60497 | 2.561 | | 0.01045 * |
|  | GenAD:Session4 | | 1.47637 | 0.63775 | 2.315 | | 0.02062 * |
|  | TreatCTSB:Session2 | | 0.05153 | 0.60969 | 0.085 | | 0.93264 |
|  | TreatCTSB:Session3 | | -0.04091 | 0.60177 | -0.068 | | 0.94579 |
|  | TreatCTSB:Session4 | | 0.23679 | 0.61103 | 0.388 | | 0.69837 |
|  | GenAD:TreatCTSB:Session2 | | -2.45644 | 0.85187 | -2.884 | | 0.00393 ** |
|  | GenAD:TreatCTSB:Session3 | | -1.02730 | 0.82343 | -1.248 | | 0.21218 |
|  | GenAD:TreatCTSB:Session4 | | -1.04578 | 0.85272 | -1.226 | | 0.22005 |
| **Diagnosis** | Residuals normality problems detected;  Quantile deviations detected; | | | | | | |
| **Other information** | Before starting the analysis, the data were corrected for the use of Beta distribution through the following procedure:  For each Session, each freezing behavior value (y) was obtaining as a frequency of the total duration of the session (i.e., 0 < y < 1), as follows:  Freezing behavior (s) / 330 (s);  Dispersion parameter for Beta family taken to be 5.7 | | | | | | |

SD = Standard Deviation; β = estimated coefficients for the fixed effect terms; SE = Standard Error; Intercept = WT-CON Session 1; Significance codes: *** 0.001; ** 0.01; * 0.05

**Table S14: Summary of the selected model for analysis of mean number of DCX^+^ cells in the dentate gyrus of the hippocampus**

| **Method** | Generalized linear model (GLM) | | | | |
| --- | --- | --- | --- | --- | --- |
| **Family distribution with link function** | Gamma ( log ) | | | | |
| **Formula** | DCX+ cells ~ Genotype * Treatment | | | | |
| **Fixed effects** |  | **β** | **SE** | **t value** | **Pr(>\|t\|)** |
|  | Intercept | 2.3023 | 0.1329 | 17.321 | < 2e-16 *** |
|  | GenAD | -0.6702 | 0.1994 | -3.362 | 0. 001886 ** |
|  | TreatCTSB | -0. 3672 | 0. 1994 | -1. 842 | 0. 074014 |
|  | GenAD:TreatCTSB | 1.0129 | 0. 2746 | 3.688 | 0. 000761 *** |
| **Diagnosis** | No diagnosis problems detected | | | | |
| **Other information** | Dispersion parameter for Gamma family taken to be 0.1766747  Null deviance: 8.4771 on 38 degrees of freedom  Residual deviance: 5.8968 on 35 degrees of freedom | | | | |

β = estimated coefficients for the fixed effect terms; SE = Standard Error; Intercept = WT-CON; Significance codes: *** 0.001; ** 0.01; * 0.05

**Table S15: Summary of the selected model for analysis of mean BrdU^+^ cell number in the dentate gyrus of the hippocampus**

| **Method** | Generalized linear model (GLM) | | | | |
| --- | --- | --- | --- | --- | --- |
| **Family distribution with link function** | Gamma ( log ) | | | | |
| **Formula** | BrdU+ cells ~ 1 | | | | |
| **Fixed effects** |  | **β** | **SE** | **t value** | **Pr(>\|t\|)** |
|  | Intercept | 0.1248 | 0.1502 | 0.831 | 0.411 |
| **Diagnosis** | Outlier detected;  No other diagnosis problems detected | | | | |
| **Other information** | Dispersion parameter for Gamma family taken to be 0.9019985  Null deviance: 110.23 on 39 degrees of freedom  Residual deviance: 110.23 on 39 degrees of freedom  Estimated Marginal Means and their respective 95% Confidence Intervals  (values are presented as number of cells)  Population = 1.13; 0.84 – 1.54 | | | | |

β = estimated coefficients for the fixed effect terms; SE = Standard Error; Intercept = Overall; Significance codes: *** 0.001; ** 0.01; * 0.05

**Table S16: Summary of the Spearman's rank correlation rho test for analysis of correlation between DCX^+^ cells and BrdU^+^ cells**

| **Method** | Spearman's rank correlation rho | | | | |
| --- | --- | --- | --- | --- | --- |
| **Alternative Hypothesis** | H1 = true rho (ρ) is not equal to 0 | | | | |
| **Output** |  | **S value** | **N** | **rho (ρ)** | **p-value** |
|  | Population | 5131 | 39 | 0.481 | 0.0020 * |
|  | WT-CON | 64 | 10 | 0.612 | 0.0665 |
|  | WT-CTSB | 52.12 | 8 | 0.380 | 0.3538 |
|  | AD-CON | 88.024 | 8 | -0.048 | 0.9103 |
|  | AD-CTSB | 126 | 13 | 0.654 | 0.0183 * |

N= sample size; rho (ρ) = Spearman correlation coefficient; Significance codes: *** 0.001; ** 0.01; * 0.05

**Table S17: Summary of the selected model for analysis of density of ThioS^+^ labeling in the cortex**

| **Method** | Generalized linear model (GLM) | | | | |
| --- | --- | --- | --- | --- | --- |
| **Family distribution with link function** | Beta ( logit ) | | | | |
| **Formula** | Density of ThioS ~ 1 | | | | |
| **Fixed effects** |  | **β** | **SE** | **z value** | **Pr(>\|z\|)** |
|  | Intercept | -3.6698 | 0.1639 | -22.38 | <2e-16 *** |
| **Diagnosis** | No diagnosis problems detected | | | | |
| **Other information** | Before starting the analysis, the data were corrected for the use of Beta distribution through the following procedure:  Each percentage of ThioS+ area value (y) was obtaining as a frequency value (i.e., 0 < y < 1), as follows:  ThioS+ area (%) / 100;  Dispersion parameter for Beta family taken to be 79.8  Estimated Marginal Means and their respective 95% Confidence Intervals  (values are presented as % of area)  Population = 2.48; 1.77 – 3.48 | | | | |

β = estimated coefficients for the fixed effect terms; SE = Standard Error; Intercept = Overall; Significance codes: *** 0.001; ** 0.01; * 0.05

**Table S18: Summary of the selected model for analysis of density of ThioS^+^ labeling in the hippocampus**

| **Method** | Generalized linear model (GLM) | | | | |
| --- | --- | --- | --- | --- | --- |
| **Family distribution with link function** | Beta ( logit ) | | | | |
| **Formula** | Density of ThioS ~ 1 | | | | |
| **Fixed effects** |  | **β** | **SE** | **z value** | **Pr(>\|z\|)** |
|  | Intercept | -4.0747 | 0.2058 | -19.8 | <2e-16 *** |
| **Diagnosis** | No diagnosis problems detected | | | | |
| **Other information** | Before starting the analysis, the data were corrected for the use of Beta distribution through the following procedure:  Each percentage of ThioS+ area value (y) was obtaining as a frequency value (i.e., 0 < y < 1), as follows:  ThioS+ area (%) / 100;  Dispersion parameter for Beta family taken to be 74.6  Estimated Marginal Means and their respective 95% Confidence Intervals  (values are presented as % of area)  Population = 1.67; 1.09 – 2.56 | | | | |

β = estimated coefficients for the fixed effect terms; SE = Standard Error; Intercept = Overall; Significance codes: *** 0.001; ** 0.01; * 0.05

**Table S19: Summary of the selected model for analysis of ThioS^+^ plaque number in the cortex**

| **Method** | Generalized linear model (GLM) | | | | |
| --- | --- | --- | --- | --- | --- |
| **Family distribution with link function** | Gamma ( log ) | | | | |
| **Formula** | Number of ThioS ~ 1 | | | | |
| **Fixed effects** |  | **β** | **SE** | **z value** | **Pr(>\|z\|)** |
|  | Intercept | 4.0934 | 0.1336 | 30.63 | <2e-16 *** |
| **Diagnosis** | No diagnosis problems detected | | | | |
| **Other information** | Dispersion parameter for Gamma family taken to be 0.339  Estimated Marginal Means and their respective 95% Confidence Intervals  (values are presented as % of area)  Population = 59.9; 45.2– 79.5 | | | | |

β = estimated coefficients for the fixed effect terms; SE = Standard Error; Intercept = Overall; Significance codes: *** 0.001; ** 0.01; * 0.05

**Table S20: Summary of the selected model for analysis of ThioS^+^ plaque number in the hippocampus**

| **Method** | Generalized linear model (GLM) | | | | |
| --- | --- | --- | --- | --- | --- |
| **Family distribution with link function** | Gamma ( log ) | | | | |
| **Formula** | Number of ThioS ~ 1 | | | | |
| **Fixed effects** |  | **β** | **SE** | **z value** | **Pr(>\|z\|)** |
|  | Intercept | 3.8516 | 0.2132 | 18.06 | <2e-16 *** |
| **Diagnosis** | No diagnosis problems detected | | | | |
| **Other information** | Dispersion parameter for Gamma family taken to be 0.864  Estimated Marginal Means and their respective 95% Confidence Intervals  (values are presented as % of area)  Population = 47.1; 30.0 – 73.8 | | | | |

β = estimated coefficients for the fixed effect terms; SE = Standard Error; Intercept = Overall; Significance codes: *** 0.001; ** 0.01; * 0.05

**Table S21: Summary of the selected model for analysis of density of Iba1^+^ labeling in the hippocampus**

| **Method** | Generalized linear model (GLM) | | | | |
| --- | --- | --- | --- | --- | --- |
| **Family distribution with link function** | Beta ( logit ) | | | | |
| **Formula** | Density of Iba1 ~ Genotype | | | | |
| **Fixed effects** |  | **β** | **SE** | **z value** | **Pr(>\|z\|)** |
|  | Intercept | -3.4050 | 0.1130 | -30.121 | <2e-16 *** |
|  | GenAD | 0.3612 | 0.1424 | 2.536 | 0.0112 * |
| **Diagnosis** | No diagnosis problems detected | | | | |
| **Other information** | Before starting the analysis, the data were corrected for the use of Beta distribution through the following procedure:  Each percentage of Iba1+ area value (y) was obtaining as a frequency value (i.e., 0 < y < 1), as follows:  Iba1+ area (%) / 100;  Dispersion parameter for Beta family taken to be 131  Estimated Marginal Means and their respective 95% Confidence Intervals  (values are presented as % of area)  WT = 3.21; 2.57 – 4.01  AD = 4.55; 3.80 – 5.43 | | | | |

β = estimated coefficients for the fixed effect terms; SE = Standard Error; Intercept = WT; Significance codes: *** 0.001; ** 0.01; * 0.05

**Table S22: Summary of the selected model for analysis of density of GFAP^+^ labeling in the hippocampus**

| **Method** | Generalized linear model (GLM) | | | | |
| --- | --- | --- | --- | --- | --- |
| **Family distribution with link function** | Beta ( logit ) | | | | |
| **Formula** | Density of GFAP ~ 1 | | | | |
| **Fixed effects** |  | **β** | **SE** | **z value** | **Pr(>\|z\|)** |
|  | Intercept | -1.19702 | 0.07697 | -15.55 | <2e-16 *** |
| **Diagnosis** | No diagnosis problems detected | | | | |
| **Other information** | Before starting the analysis, the data were corrected for the use of Beta distribution through the following procedure:  Each percentage of GFAP+ area value (y) was obtaining as a frequency value (i.e., 0 < y < 1), as follows:  GFAP+ area (%) / 100;  Dispersion parameter for Beta family taken to be 23.9  Estimated Marginal Means and their respective 95% Confidence Intervals  (values are presented as % of area)  Population = 23.2; 20.5 – 26.1 | | | | |

β = estimated coefficients for the fixed effect terms; SE = Standard Error; Intercept = Overall; Significance codes: *** 0.001; ** 0.01; * 0.05
